# Dark-induced inactivation of the carbon assimilation process requires a water–water cycle–driven oxidative burst

**DOI:** 10.64898/2026.01.10.698556

**Authors:** Matanel Hipsch, Moshe Steinberg, Itay David, Shai Nouriel, Shilo Rosenwasser

## Abstract

Chloroplast metabolism is constantly fine-tuned by light availability through the perception and transmission of reductive and oxidative signals that activate or deactivate distinct metabolic enzymes. The reducing power originating from the photosynthetic electron transport chain has been shown to fuel the redox regulatory network, linking electron transport to the reductive activation of photosynthetic enzymes. However, the source of the oxidizing equivalents required to reverse photosynthetic enzyme activation and drive them toward an oxidized, inactive state has not yet been experimentally demonstrated. Here, we resolve redox dynamics associated with carbon assimilation inactivation by combining time-resolved redox imaging during the light-to-dark transition (LDT) with gas-exchange–based measurements. Dark-induced inactivation of carbon assimilation proved oxygen-dependent and coincided with an oxygen-dependent oxidative burst triggered during the LDT. This oxidative burst was suppressed under conditions that blocked electron transport to PSI or in plants in which PSI was photoinactivated. Notably, *pgr5* and *pgrl1ab* mutants exhibited attenuated oxidative bursts and suppressed LDT-associated carbon assimilation inactivation, demonstrating that PGR5/PGRL1-dependent activity is required to generate the oxidative burst that drives CBC inactivation during LDT. These results establish a direct mechanistic link between oxygen- and PSI-dependent oxidative bursts and the inhibition of photosynthesis and mark the water–water cycle (WWC) as the primary source of the transient accumulation of oxidative equivalents that drive inactivation of Calvin–Benson cycle enzymes in darkness.

## Introduction

The daily plant metabolism cycle shifts from photoautotrophic, relying on active photosynthesis during the day, to heterotrophic, relying on starch metabolism for energy at night. Transitions between these two metabolic states can also intermittently occur throughout the day in response to dynamic, fluctuating light conditions. Maintaining homeostasis under such dynamic conditions necessitates regulatory mechanisms that enable the rapid activation and inactivation of metabolic enzymes.

Redox regulation mediated by thiol–disulfide exchange and driven by the transmission of reductive and oxidative signals from the photosynthetic electron transport chain (PETC) to target proteins is a key mechanism that enables chloroplast metabolism to rapidly adjust to changes in light intensity (Buchanan, 2014; Buchanan & Balmer, 2005; Schürmann & Buchanan, 2008). Typically, the activation of redox-regulated enzymes depends on reductive signals that reduce disulfide bonds, whereas the reception of oxidative signals results in disulfide bond formation and enzyme inactivation (Buchanan, 1981).

Reductive activation signals are mediated by the flux of electrons from ferredoxin (Fd) to thioredoxins (Trxs) via Fd-Trx reductase (FTR), or through Fd–NADP⁺ reductase (FNR) and NADPH-dependent Trx reductase C (NTRC) (Buchanan & Balmer, 2005; Michalska et al., 2009; Serrato et al., 2004; Yoshida et al., 2022). This redox cascade initiates carbon assimilation in response to light by activating key Calvin–Benson cycle (CBC) enzymes, such as fructose-1,6-bisphosphatase (FBPase), sedoheptulose-1,7-bisphosphatase (SBPase), phosphoribulokinase (PRK) and glyceraldehyde-3-phosphate dehydrogenase (GAPDH) (Buchanan et al., 2012; Buchanan & Balmer, 2005; Motohashi et al., 2001; Souza, 2025).

Oxidative inactivation signals are mediated by high-midpoint-potential atypical Trxs and 2-Cys peroxiredoxins (Prxs), which catalyze the transfer of reducing equivalents from target proteins to H_2_O_2_, thereby shutting down protein activity (Dangoor et al., 2012; Eliyahu et al., 2015; Ojeda et al., 2018; Vaseghi et al., 2018; Yoshida et al., 2018). This oxidizing activity relies on the buildup of an oxidizing equivalent that draws reducing power from redox-regulated enzymes. However, while light-induced generation of reduced Fd underlies reductive activity, the sources of H₂O₂ required as electron acceptors for protein inactivation in the dark, as well as the mechanisms underlying its formation, remain poorly defined.

Reactive oxygen species (ROS) production within the photosynthetic apparatus has been proposed to occur at multiple sites (Fantuzzi et al., 2022; Foyer & Hanke, 2022; Lee & Kim, 2024), with the donation of electrons from Photosystem I (PSI) to molecular oxygen in the Mehler reaction being the major route for stromal superoxide formation (Mehler, 1951). The production of superoxide initiates a set of reactions known as the water–water cycle (WWC) (Asada, 1999; Badger, 1985), in which superoxide (O_2−_) is initially dismutated to molecular oxygen (O_2_) and H_2_O_2_ in a reaction catalyzed by superoxide dismutase (SOD) (Asada et al., 1974; Mehler, 1951). Subsequent reduction of H₂O₂ to water is catalyzed by ascorbate peroxidase (APX), with the regeneration of ascorbate achieved through the ascorbate-glutathione (GSH) cycle (Foyer & Noctor, 2011; Terai et al., 2020). Alternatively, H₂O₂ detoxification to water can be mediated by 2-Cys-Prx (Dietz et al., 2006; König et al., 2002; Vogel et al., 2014). Collectively, in the WWC, electrons originating from water at Photosystem II (PSII) reduce O₂ back to water in the stroma.

Within PSI, the photoreduction of O_2_ to superoxide has been suggested to occur at FeS_x_ and FeS_A/B_ clusters, which transfer electrons from the PSI reaction center to Fd (Asada, 1999). Accordingly, high-light(HL)-induced damage to FeS_A/B_ resulted in low superoxide levels in Arabidopsis WT plants (Tiwari et al., 2024). When the FeSₓ clusters are damaged, as occurs in HL-–exposed proton gradient regulator 5 mutant plants *(pgr5)*, which lack key mechanisms that protect PSI from photoinhibition, low levels of superoxide are generated even in the presence of methyl viologen (MV), which induces superoxide production in WT (Tiwari et al., 2024). Similarly, *pgr5* plants exhibited decreased production of superoxide and H_2_O_2_, increased antioxidant capacity (Suorsa et al., 2012) and failed to induce chloroplastic glutathione redox potential (*E_GSH_*) oxidation under fluctuating light conditions (Haber et al., 2021), underscoring the importance of functional PSI in regulating chloroplastic ROS metabolism.

Several physiological functions have been attributed to WWC (Foyer & Kunert, 2024), including its role as an electron sink to prevent over-reduction of the PETC, in maintaining PSI redox balance when the supply of reducing equivalents exceeds their consumption by downstream metabolism (Miyake et al., 2006; Park et al., 1996), in promoting non-photochemical quenching (NPQ) by contributing to ΔpH formation across the thylakoid membrane (Schreiber & Neubauer, 1990), and in adjusting the ATP/NADPH ratio to support downstream metabolic demands (Asada, 2000; Miyake, 2010; Sun et al., 2020). Although these functions underscore the benefits of routing electrons via the WWC, they do not assign any functional role to ROS; rather, ROS are regarded as incidental by-products whose levels are constrained by the antioxidant network. However, the limited capacity of the WWC in C₃ plants, along with the lack of evidence for its involvement in activating photoprotective mechanisms, raises questions about its role as an alternative sink or NPQ induction, and instead suggests that ROS generated via the WWC functions in signaling (Badger et al., 2000; Driever & Baker, 2011; Exposito-Rodriguez et al., 2017). Additionally, a regulatory role for WWC-derived ROS may explain its evolutionary conservation, while mechanisms allowing direct PSI-to-O₂ electron transfer without ROS production, mediated by flavodiiron proteins, were lost in flowering plants (Ilík et al., 2017; Zhang et al., 2009).

A key regulatory role for chloroplastic ROS is further supported by the measurable ROS levels detected in chloroplasts despite their highly efficient antioxidant system, as indicated by the light-induced oxidation of H₂O₂ biosensors such as Hyper (Exposito-Rodriguez et al., 2017) and roGFP2-PrxΔC_R_ (Hipsch et al., 2025; Lampl et al., 2022). The recent elucidation of the 2-Cys Prx-dependent oxidative pathway (Dangoor et al., 2012; Eliyahu et al., 2015; Ojeda et al., 2018; Vaseghi et al., 2018; Yoshida et al., 2018) raises the possibility that WWC-derived H₂O₂ production serves as a sink for electrons withdrawn from redox-regulated proteins. Yet, direct evidence linking WWC activity to the physiological inactivation of photosynthesis has been missing.

Here, by combining redox imaging with quantitative, non-destructive assessments of CBC inactivation in Arabidopsis and potato under pharmacological, photoinhibitory and genetic perturbations of photosynthetic electron chain, we demonstrated the link between PSI-dependent O₂ photoreduction, PGR5/ PGRL1 activity, and inactivation of carbon assimilation, demonstrating the role of WWC and PGR5-PGRL1 complex in facilitating the condition required for oxidative inactivation of photosynthetic enzymes.

## Results

### Oxygen is required for the inactivation of carbon assimilation during the transition from light to dark

Disruption of the oxidative regulatory pathways results in retention of the CBC enzymes in a reduced state after the transition to darkness (Ojeda et al., 2018; Vaseghi et al., 2018; Yoshida et al., 2018; Doron et al., 2025). Such inhibition of dark-induced oxidation triggers a faster-than-wild-type (WT) photosynthetic induction upon reillumination, as detected in Arabidopsis mutants lacking 2-Cys Prx A and B (*2cpab*) (Lampl et al., 2022), suggesting that the redox state of photosynthetic enzymes governs CO_2_ assimilation rates at the beginning of the induction phase (Sassenrath-Cole & Pearcy, 1994). As such, measuring the rate of CO₂ assimilation upon reillumination of dark-adapted plants enables exploration of the oxidative inactivation process induced during the transition to darkness (Fig. 1a). Indeed, measuring CO₂ assimilation during photosynthetic induction in plants exposed to 1-12 min of dark revealed a dark-duration-dependent photosynthesis slowdown; the longer the duration of the dark period, the more attenuated the slope of photosynthetic recovery upon transition to light was, demonstrating that the inactivation is a continuous process (Fig. 1b).

**Figure 1:**
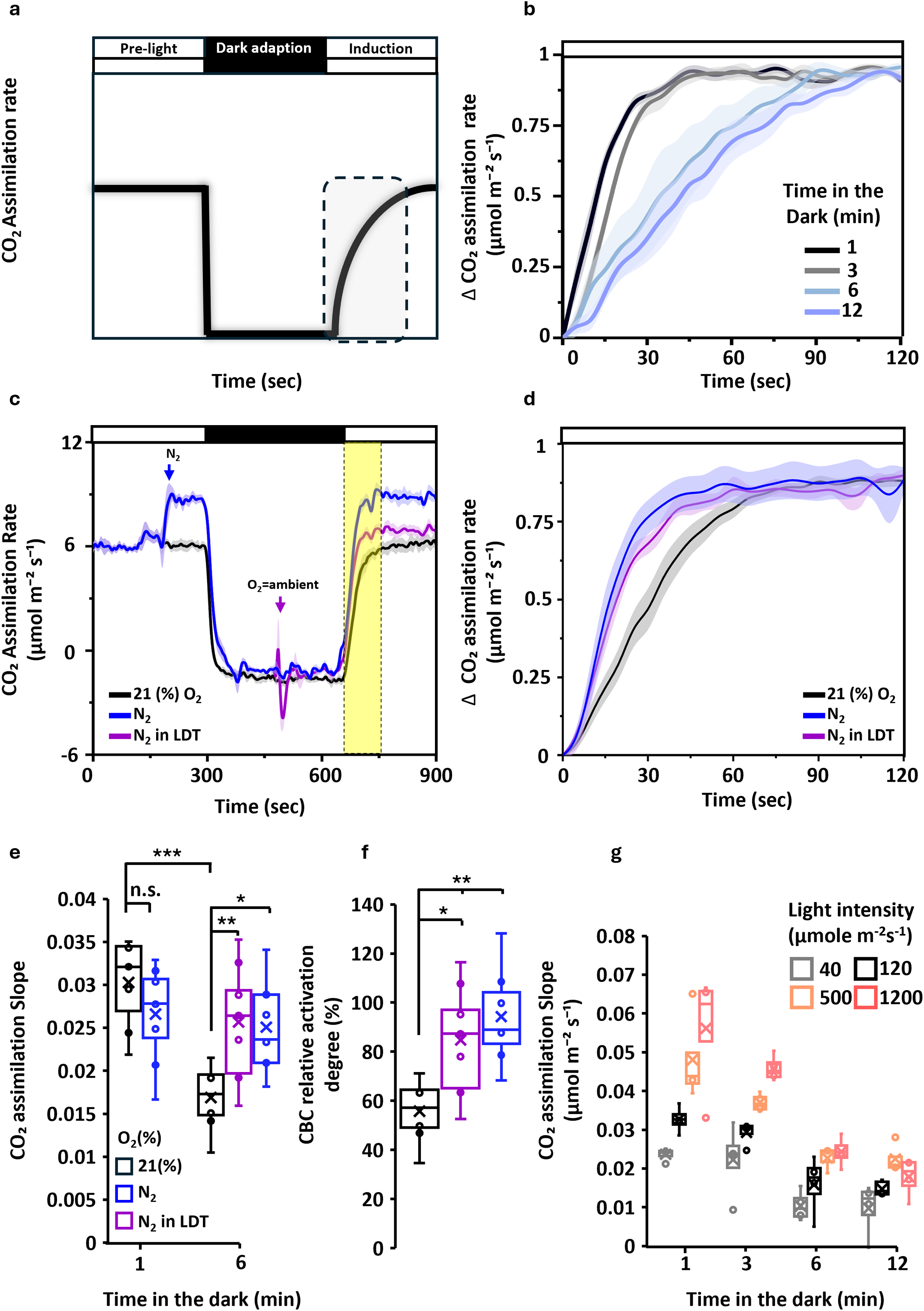
Oxygen availability and light intensity shape the kinetics of carbon assimilation inactivation during light-to-dark transitions. a) Schematic illustration of the methodology used to assess carbon assimilation inactivation. Inactivation was evaluated by measuring the slope of CO₂-assimilation induction upon reillumination after a dark period that was preceded by a defined light treatment. b) Reactivation of carbon assimilation in intact potato leaves exposed to 120 µmol m⁻² s⁻¹ light after dark periods of varying durations (1, 3, 6 and 12 min). Data are shown as mean ± SE (n = 4), with shaded areas indicating ±SE. c) Carbon assimilation rate was measured during the light–dark–light treatment shown in Fig. 1a, under 21% O₂, continuous N₂ exposure (initiated after 3 min of light), or N₂ during light to dark transitions (N₂ in LDT). The region marked in yellow indicates the interval over which slopes were calculated (see panel e). Data are expressed as mean ± SE (n = 6-7), with shaded regions representing ± SE. Arrows show the timing of N_2_ insertion (blue, 3 min) or depletion (purple, 3 min in the dark) from the system. d) Normalized carbon assimilation rate during photosynthesis induction, in potato leaves subjected to the treatment described in (c). Data are expressed as mean ± SE (n = 6-7). e) Box plots showing the CO₂-assimilation induction slopes of potato leaves subjected to the treatment described in (d), following dark-inactivation periods of either 1 or 6 min. Values are presented as mean ± SE (n = 6–9). f) Box plots showing the CBC inactivation degree, calculated as (See method). Values are presented as mean (n = 6–9). g) Box plots showing the CO₂-assimilation induction slopes of potato plants exposed to 40, 120, 500 or 1200 µmol m⁻² s⁻¹ light intensities, followed by dark periods of 1–12 min, and 5 min of reactivation at 120 µmol m⁻² s⁻¹. Values are shown as mean ± SE (n = 4–5). Statistical significance was assessed for panels (e and f) using a two-tailed Student’s t-test and is marked as p < 0.05 (*), < 0.01 (**), < 0.001 (***), while "n.s." denotes non-significant differences.

To test our hypothesis that Mehler reaction-derived H₂O₂ production is essential for inactivating CBC-related enzymes during the light-dark transition (LDT), the effect of limited O₂ availability on protein inactivation in the dark was assessed. More specifically, CO_2_ assimilation was examined during photosynthetic induction in leaves dark-adapted for 1 and 6 min and incubated in N₂ to limit oxygen reduction and H_2_O_2_ production. CO_2_ assimilation measurements were limited to 6 minutes of dark duration to exclude possible effects of stomatal closure, which may influence photosynthetic induction during extended dark adaptation. As expected, stomata remained unresponsive for at least 6 min in darkness, consistent with their slower kinetics relative to redox changes (Fig. Supp. 1, McAusland et al., 2016). Under ambient air, the normalized induction slope declined from 0.3 after 1-min dark adaptation to 0.17 after 6 min, representing a 45% decrease in the activation degree of CBC-related enzymes (See methods, Fig. 1c-f). In N₂-treated plants, slopes declined from 0.265 to 0.25, reflecting a 6% drop in enzyme activation state (Fig. 1c–f), indicating that O₂ plays a role in the dark-induced inactivation. To rule out the possible effect of photorespiration on the induction dynamics, the experiment was repeated in plants exposed to N₂ during the first light phase and during the transition to dark, while exposed to air before the light was turned on. A markedly steeper induction slope of 0.25 was measured in plants exposed to N₂ compared with those kept under ambient O₂ (0.17) (Fig. 1c–f). Compared with plants exposed to N₂ throughout the experiment, activation decreased by 16% in plants exposed to N₂ during the transition and by 44% in plants exposed to ambient oxygen conditions. Together, these results demonstrate that O₂ availability during LDT is critical for CBC inactivation.

To further investigate the inactivation dynamics in plants exposed to different light intensities, plants were pre-illuminated with 40, 120, 500, or 1200 µmol photons m²s¹, followed by a dark period and re-exposure to 120 µmol m-²s-¹ (Fig. 1f). After 1 min in darkness, assimilation slopes were 0.56, 0.48, 0.32 and 0.23 at 1200, 500, 120, and 40 µmol m²s¹, respectively (Fig. 1g), indicating that HL intensity results in a higher activation state of CBC-related enzymes. After 6 min of darkness, the assimilation rates across all treatments converged, indicating that the CBC-related enzymes reached a common inactivation state, regardless of prior light exposure. This inactivation dynamic is consistent with the observation that FBPase, GAPDH and Rubisco activase (RCA) enzymes reached 50-80% oxidation after 5 min in the dark (Jiménez-López et al., 2025; Ojeda et al., 2018). To conclude, these findings support a model in which light-dependent H₂O₂ production serves as the primary electron acceptor for the CBC enzyme inactivation in darkness, with both light intensity and dark duration shaping the dynamics of inactivation.

### Oxygen-dependent oxidative peak in stromal *E_GSH_* during light-to-dark transition

We reasoned that H_2_O_2_ bursts originating from photosynthetic machinery during LDT can drain electrons from GSH for H_2_O_2_ detoxification via the ascorbate-GSH cycle, thereby increasing the redox potential of *E_GSH_*. Therefore, Arabidopsis and potato plants expressing the chl-roGFP2 biosensors were used to monitor *E_GSH_* during LDT using *in vivo* imaging (Haber et al., 2021; Hipsch et al., 2021). Chloroplast localization of the probe was confirmed by confocal microscopy and the dynamic range of probe’s (R₄₀₅/₄₆₅ dynamic ranges of 3.7–5.97 for the tested lines) was demonstrated using H_2_O_2_ and dithiothreitol (DTT) treatments (Supp. Fig. 2). Following the transfer of biosensor lines grown under standard light conditions (GL, 120 µmol m² s¹) to darkness, the degree of roGFP2 oxidation peaked at 12 min and then gradually returned to steady-state levels (Fig. 2a, c). The LDT-induced increase in roGFP2 oxidation degree (OxD), which measured 25±0.8% (SE) and 42±2.2% (SE) for potato and Arabidopsis, respectively, indicated a 10-20 mV increase in the redox potential of *E_GSH_*. This oxidative peak was consistent across whole plants, detached leaves (DL) and leaf discs (LD) (Fig. 2a-c). Similar oxidation trends were observed in Arabidopsis-chl-Grx1-roGFP2, in which Grx1 is fused to roGFP2 (Marty et al., 2009). As Grxs mediate equilibration between roGFP2 and the GSH pool, this result suggests that sufficient endogenous Grxs are available to drive the oxidation and reduction of roGFP2. Notably, roGFP2 targeted to mitochondria and cytosol showed no changes during LDT, confirming that this oxidation is specific to the chloroplast stroma (Supp. Fig. 3). In addition, LDT chl-roGFP2 oxidation was halted upon exposure to various probe inhibitors (Supp. Fig. 4). These results align with prior studies demonstrating chloroplastic GSH oxidation during LDT in Arabidopsis and diatoms (Bohle, Klaus, et al., 2024; Haber et al., 2021; Mizrachi et al., 2019; Müller-Schüssele, 2024; Müller-Schüssele et al., 2021). Importantly, replacing ambient oxygen with N₂ suppressed the roGFP2 burst across all experimental setups (Fig. 2d–g and Supp. Fig. 5), suggesting that oxygen-derived H₂O₂ production induces stromal *E_GSH_* oxidation and that exploring roGFP2 oxidation dynamics can provide insights into the oxidative inactivation of redox-regulated proteins.

**Figure 2:**
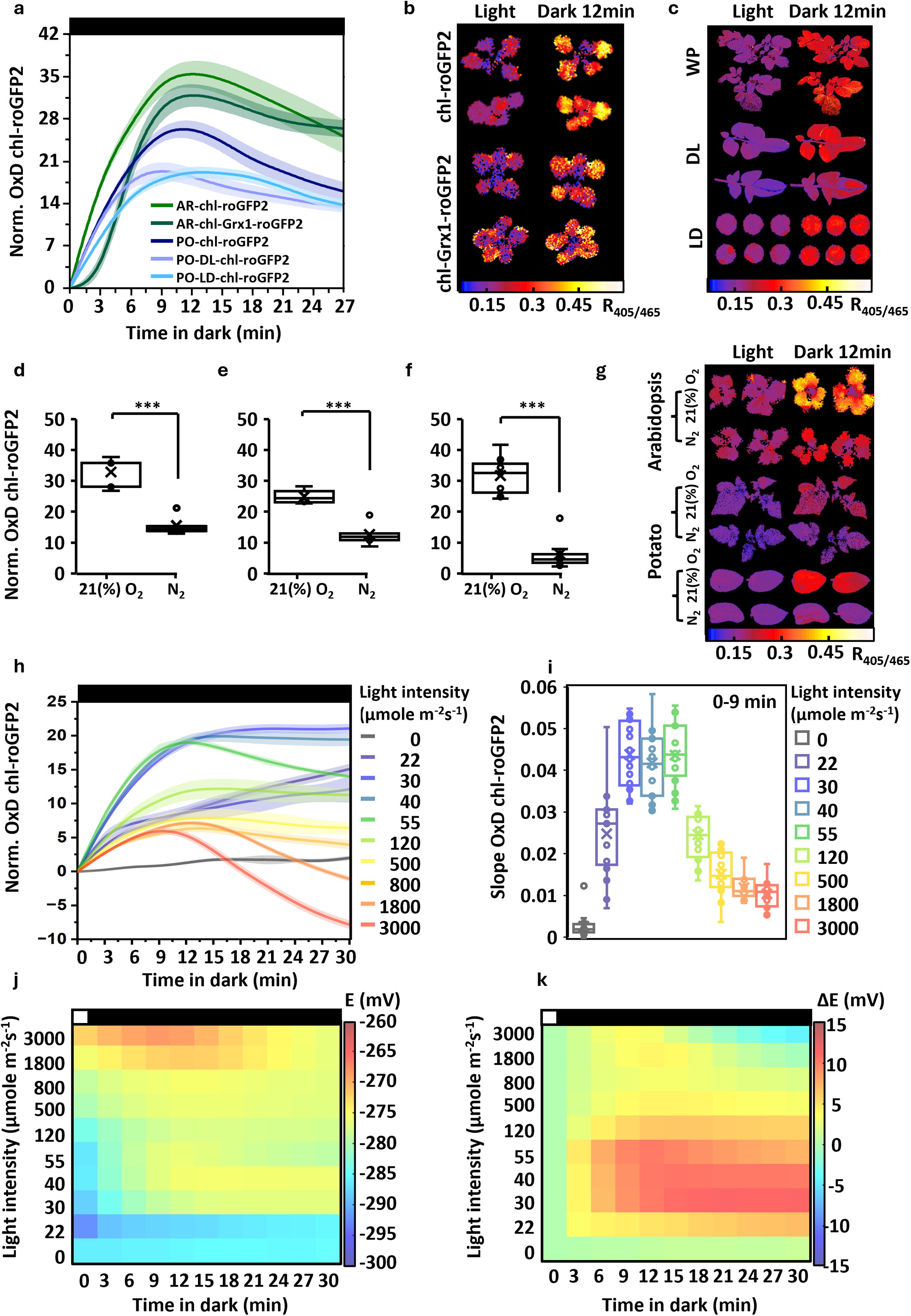
Oxidative peak in stromal *E_GSH_* during light-to-dark transition in Arabidopsis and potato. a) Changes in oxidation degree (OxD) of chl-roGFP2 in response to LDT of whole-plant (WP), detached leaves (DL) and leaf discs (LD) of Arabidopsis and potato plants. Values were normalized to OxD measured in light. For Arabidopsis, data were recorded using plants expressing the chl-roGFP2 or chl-Grx1-roGFP2 biosensors. Values represent mean ± SE (n = 3–20), with shaded areas indicating ± SE. b–c) Representative ratiometric redox images (R₄₀₅/₄₆₅) taken from potato (b) and Arabidopsis (c) at time 0 and 12 min into the LDT. d-f) Box plots showing the increase in OxD of chl-roGFP2 in *Arabidopsis* WP (d), potato WP (e), and potato DL (f), exposed to ambient O₂ or N₂ during LDT. Values were calculated as (OxD after a 12-min dark exposure) – (OxD in the light). Data are presented as mean ± SE (n = 5-9). g) Representative ratiometric redox images from the experiment described in d-f, captured at 0 and 12 min into LDT. h-i) chl-roGFP2 oxidation (h) and induction slopes (i) during LDT in plants exposed to varying light intensities (0–3000 µmol m⁻²s⁻¹) before the dark period. Slopes were calculated from changes observed during the first 0–9 min in darkness. Values represent mean ± SE (n = 18–22), with shaded areas indicating ±SE. j-k) Heat map representation of stromal glutathione redox potential (*E_GSH_*) during LDT. Panel (j) shows absolute *E_GSH_* values under light conditions and panel (k) presents the changes in *E_GSH_* between light and dark (ΔmV). Statistical significance was assessed for panels (d and e) using a two-tailed Student’s t-test and is marked as p < 0.05 (*), < 0.01 (**), < 0.001 (***), while "n.s." denotes non-significant differences.

**Figure 3:**
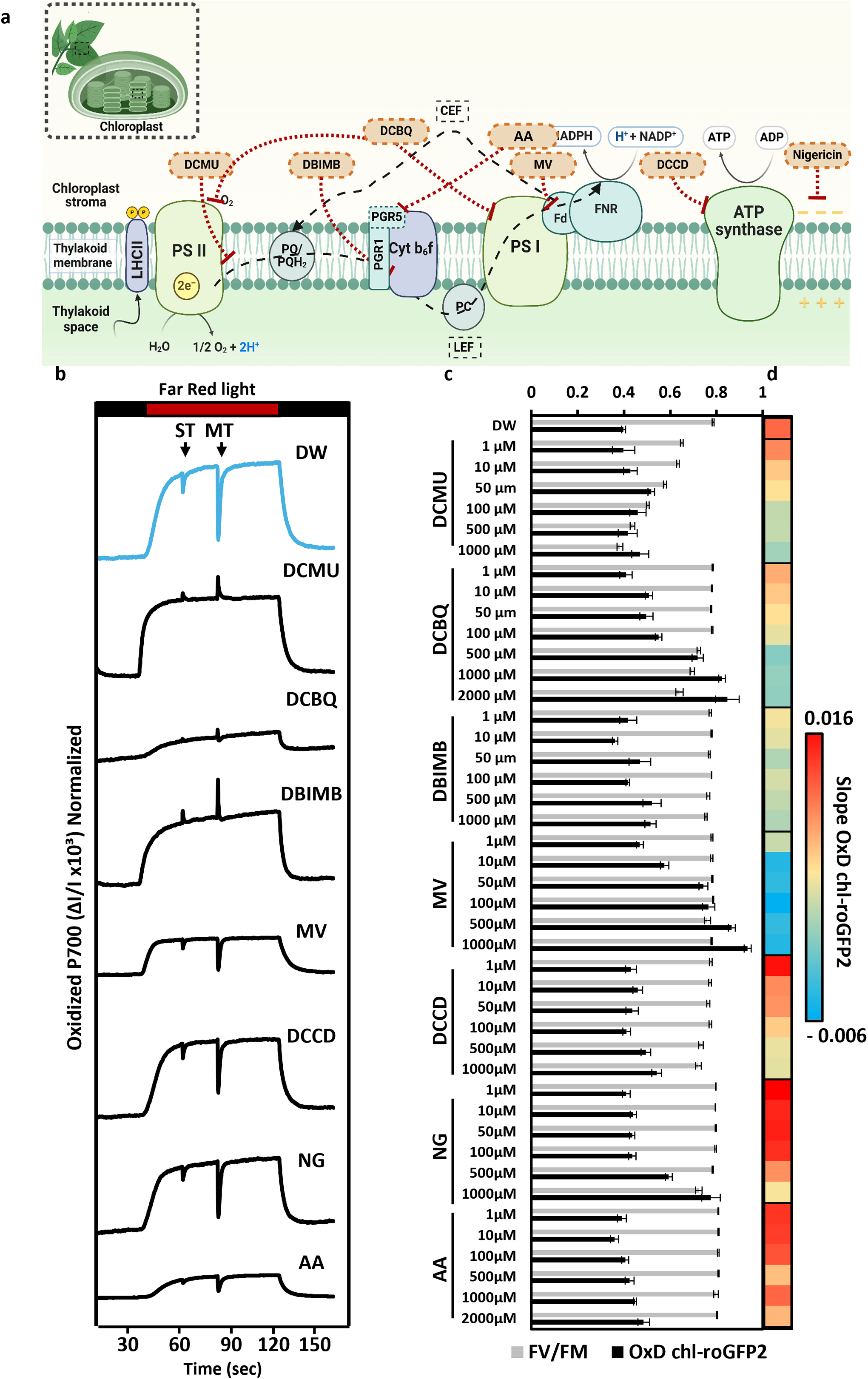
Photosynthetic electron-transport inhibitors differentially modulate the chl-roGFP2 oxidative burst during the light–dark transition. a) Schematic model of the photosynthesis apparatus, highlighting the major photosynthetic protein complexes, the inhibitors applied, and their suggested mode of action. b) P700 redox kinetics in intact leaves, showing single-turn (ST) and multiple-turn (MT) measurements, with black arrows indicating the start of each. All inhibitors were applied at 1000 µM, and data were reported as mean values from 4-5 leaflets examined per treatment. c) OxD chl-roGFP2, presented as mean ± SE (n = 10–12), and maximum quantum yield of PSII (Fv/Fm), presented as mean ± SE (n = 10–12) at illumination start (time 0). e) Heatmap presentation of the chl-roGFP2 oxidation slopes, calculated based on the change during the first 0–9 minutes in darkness. The heat maps display the average oxidation response across individual leaf discs (n = 10-12).

**Figure 4:**
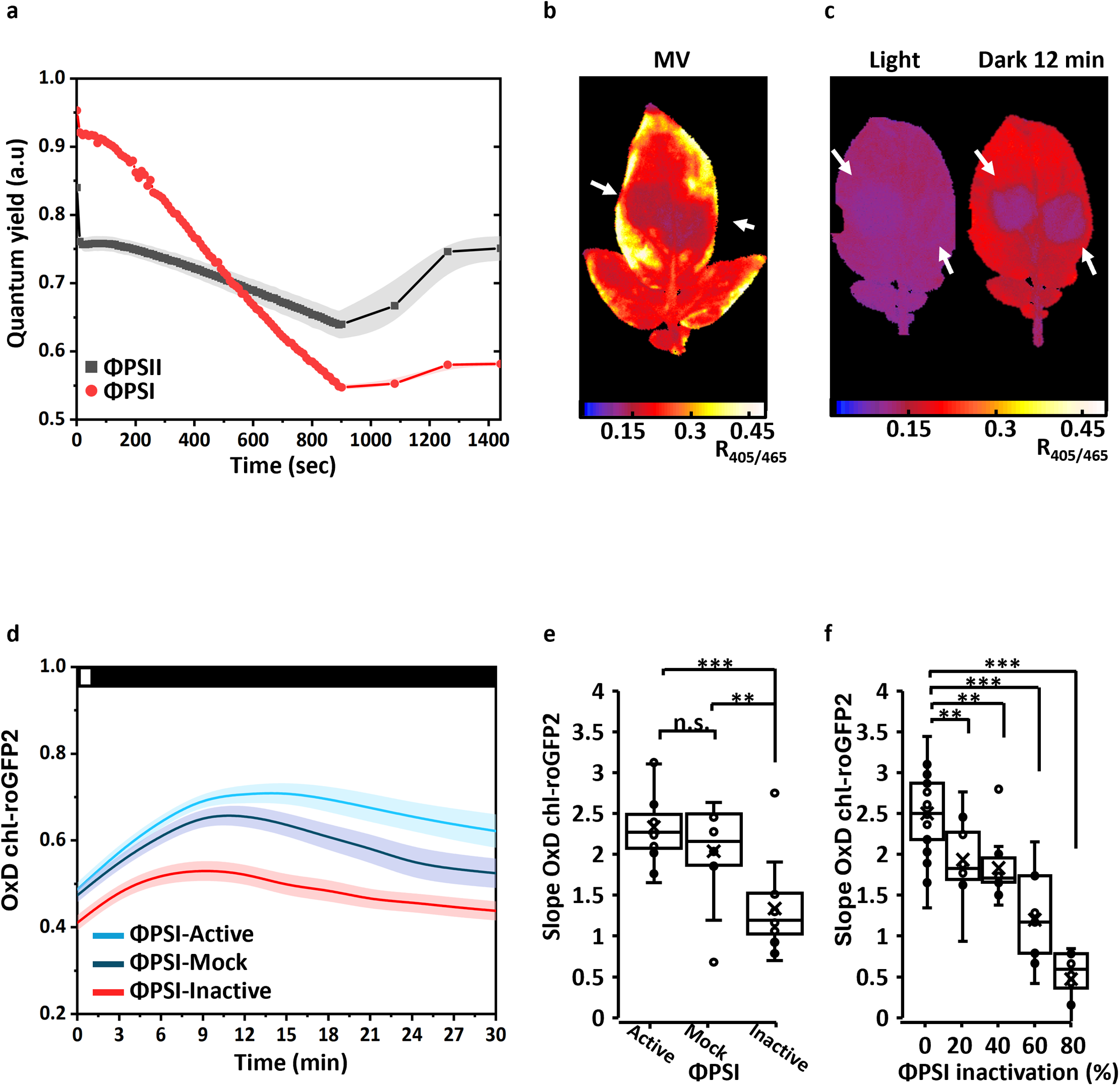
PSI activity is required for the induction of chl-roGFP2 oxidative peak during light-to-dark transition. a) Quantum yield measurements of PSII and PSI in detached leaves exposed to a repetitive saturating pulses (rSP) protocol (see Methods). Measurements were taken during the 15 min of rSP followed by a 9-min recovery phase every 3 min. Data are presented as mean ± SE (n = 4), with shaded areas indicating ±SE. b) Representative ratiometric image (R_405/465_) showing a detached leaf with inactivated PSI areas, after being soaked in 10µM methyl viologen (MV) for 1 h. White arrows indicate PSI inactivation sites. c) Representative ratiometric images (R₄₀₅/₄₆₅), taken under light conditions and after 12 min in the dark, showing PSI inactivation at two different sites. White arrows indicate PSI inactivation sites. d) Quantification of oxidation degree (OxD) of chl-roGFP2 during LDT in detached control leaves (PSI-Active), non-inactivated regions in treated leaves (PSI-Mock), and PSI-inactivated regions (PSI-Inactive). Data are shown as mean ± SE (n = 12), with shaded areas representing ±SE. e) Box plot presenting the chl-roGP2 oxidation slope derived from panel (d), calculated over the 0–9-min period following the light-to-dark transition. f) Box plot showing the dose-dependent effect of PSI inactivation (calculated based on PSI quantum yield), on chl-roGP2 oxidation slope calculated over the 0–9-min period following the light-to-dark transition (n = 9-21). Statistical significance was assessed for panels (e and f) using a two-tailed Student’s t-test and is marked as p < 0.05 (*), < 0.01 (**), < 0.001 (***), while "n.s." denotes non-significant differences.

**Figure 5:**
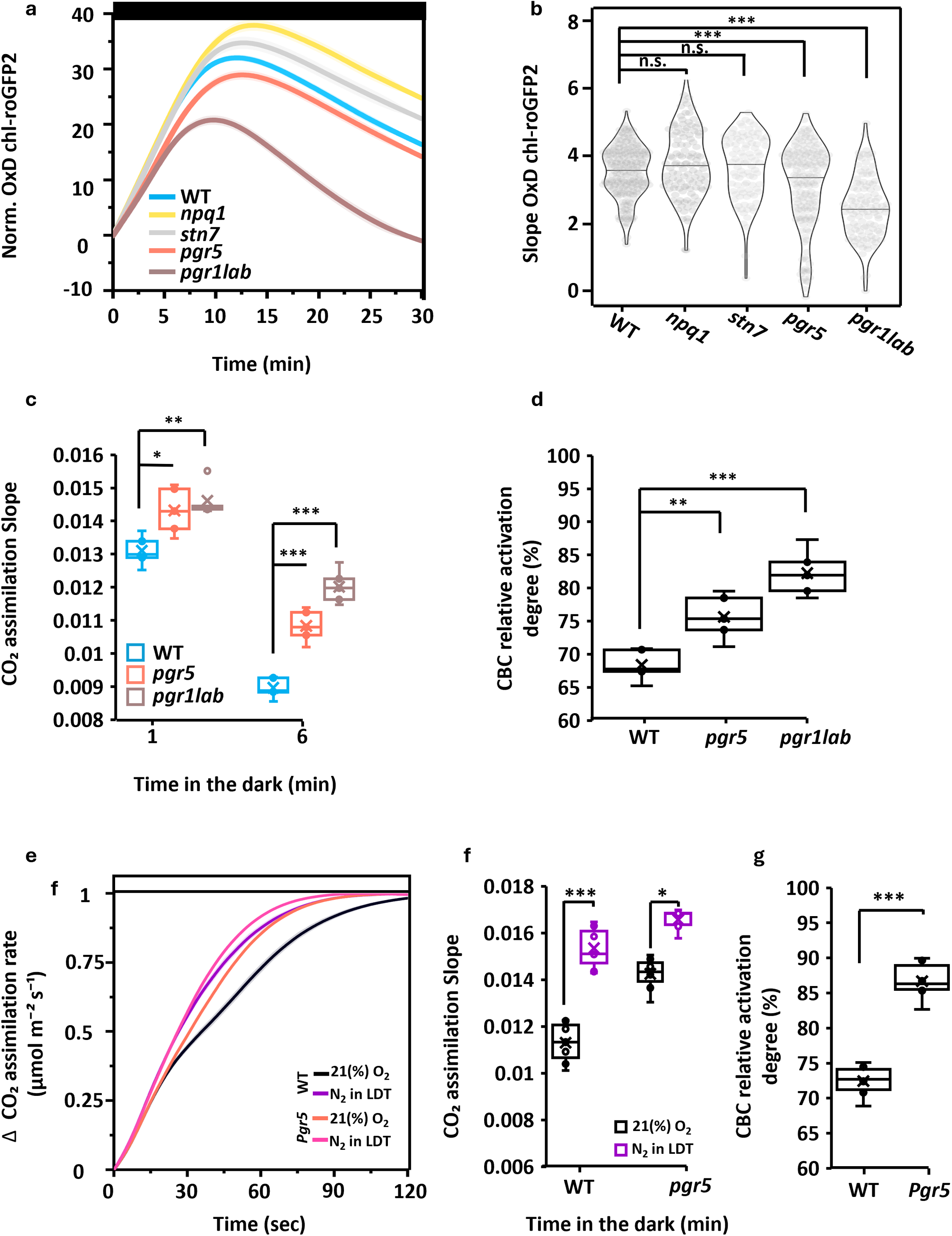
Attenuated chl-roGFP2 oxidation and suppressed inactivation of carbon assimilation during light-to-dark transition in *pgr5* and *pgrl1ab* plants. a) Changes in normalized oxidation degree (OxD) chl-roGFP2 during the LDT in *Arabidopsis* wild type (WT) and photosynthesis-related mutants (*npq1*, *stn7*, *pgr5*, *pgrl1ab*). Shaded areas represent ±SE. b) Violin plots illustrating the distribution of OxD slopes (0–9 min in darkness) for the genotypes shown in panel (a). Sample sizes per genotype: WT = 213, *npq1* = 117, *pgrl1ab* = 107, *pgr5* = 169, *stn7* = 95. c) Box plots showing normalized carbon assimilation slopes measured in Arabidopsis WT or *pgr5*, followed by dark inactivation periods of 1 or 6 min. Values are presented as mean ± SE (n = 5). d) Box plots showing the carbon assimilation activation degree of WT, *npq1*, *stn7*, *pgr5* and *pgr1lab*. Values are presented as mean (n = 5-15). e) Normalized carbon assimilation rates in Arabidopsis WT or *pgr5* plants exposed to 21% O₂, or N₂ during LDT. The graph shows the carbon assimilation rates of plants re-exposed to light following a 6-min dark period, calculated as the change from the final dark time points. Data are expressed as mean ± SE (n = 6). f) Box plots showing the CO₂ assimilation slopes measured in Arabidopsis WT or *pgr5* plants, treated with 21% O₂, or N₂ during LDT, during exposure to light followed by dark inactivation periods of 6 min. Values are presented as mean ± SE (n = 6). g) Box plots showing the carbon assimilation activation degree of Arabidopsis WT or *pgr5* plants exposed to 6-min dark inactivation and then treated with 21% O₂ or N₂ during LDT. Activation degree was calculated as. Values are presented as mean (n = 6). Statistical significance was assessed for panels (b, c, d, g, and h) using a two-tailed Student’s t-test and is marked as p < 0.05 (*), < 0.01 (**), < 0.001 (***), while "n.s." denotes non-significant differences.

### The magnitude of chloroplast E_GSH_ oxidation during LDT varies with light intensity

Given the influence of light intensity on the dynamics of carbon assimilation inactivation (Fig. 1g), subsequent experiments examined how light intensity affects the dynamics of the chl-roGFP2-related oxidative burst. chl-roGFP2 oxidation was measured in plants exposed to light intensities of 22-3000 µmol m²s¹ for 10 min, followed by a 30-min dark period (Fig. 2h-k, Supp. Fig. 6). Plants exposed to low light (LL) (up to 120 µmol m⁻² s⁻¹) displayed a significant increase of 15–20% in OxD, which saturated at 12–15 min into darkness, reflecting an *E_GSH_* shift of 15 mV from approximately –290 mV to –275 mV, with minimal subsequent reduction. Conversely, HL-exposed plants (500–3000 µmol m⁻² s⁻¹) exhibited a notably lower chl-roGFP2 oxidation peak, ranging from 6% to 7% OxD, which saturated earlier, at 6–9 min into darkness, reflecting an *E_GSH_* shift of 10 mV (Fig. 2d). Moreover, unlike LL conditions, HL-treated plants displayed, following the burst, a rapid reduction to baseline or below (at higher light 1800-3000 µmol m²s¹) within 30 min of darkness (Fig. 2j-k). These differences were reflected in changes in chl-roGFP2 oxidation slopes across light intensities, forming a bell-shaped curve (Fig. 2i). Notably, the magnitude of the oxidative burst was not affected by the time of day, as shown in experiments in which plants were monitored for 12 h under GL or exposed to a gradual light increase, mimicking natural light conditions (Supp. Fig. 7). Taken together, these results demonstrate that the dynamics of chl-roGFP2 oxidation during LDT are regulated by the light intensity experienced before the transition to dark.

### PSI activity mediates chl-roGFP2 oxidation during the transition to darkness

To further link the chloroplast roGFP2 oxidation peak during LDT to photosynthetic activity during the preceding light phase, potato plants expressing chl-roGFP2 were treated with varying concentrations of 3-(3,4-dichlorophenyl)-1,1-dimethylurea (DCMU) which inhibits PQ pool reduction by blocking the Q_B_ site in PSII (Barry et al., 1990), 6-dichloro-1,4-benzoquinone (DCBQ), a redox mediator that extracts electrons from PSI and PSII (Graan & Ort, 1986), dibromo-6-isopropyl-3-methyl-1,4-benzoquinone (DBIMB), an inhibitor of the Qo site, inhibiting PQ oxidation by Cyt-b6f, thus blocking photosynthetic electron transfer between PSII and PSI (Trebst et al., 1970), methyl viologen (MV), a catalyst for superoxide generation at the PSI acceptor side (Bus & Gibson, 1984), dicyclohexylcarbodiimide (DCCD), an inhibitor of the F0F1-ATP synthase (Jahns & Junge, 1989; Lam et al., 2024), nigericin (NG), a protonophore that collapses the proton gradient (ΔpH) across the thylakoid membrane, thereby relaxing NPQ and accelerating photoinhibition in PSII (Shavit et al., 1968) and antimycin A (AA), which blocks the antimycin A-sensitive Fd-dependent CEF(Tagawa et al., 1963) (Fig. 3, a). Significant chl-roGFP2 oxidation was observed in DCBQ, MV and NG-treated plants during the light phase, while DCMU, DBMIB, DCCD and AA induced minimal oxidation compared to Distilled Water (DW). The maximum quantum yield of PSII (Fv/Fm) values remained largely stable (∼0.8–0.75) across treatments, except for DCMU, which reduced Fv/Fm to 0.38 at 1000 µM (Fig. 3c). PETC-targeting inhibitors DCMU, DBMIB, and DCBQ, fully blocked P700 reduction under both single-turnover (ST) and multiple-turnover (MT) pulses, confirming complete inhibition of electron flow from PSII to PSI (Fig. 3, b) (Klughammer & Schreiber, 2016; Tiwari et al., 2016). These treatments also strongly suppressed the LDT oxidative burst (Fig. 3d). In contrast, moderate inhibition of the LDT oxidative response was observed following AA and NG treatments, which do not directly affect electron transfer to PSI (Fig. 3b & d). MV-induced probe oxidation in the light made it challenging to observe redox changes during LDT; nevertheless, the suppression of LDT oxidative burst was noticeable in response to 1 µM (Fig. 3C). These findings suggest that the chl-roGFP2 oxidative burst arises from within the PETC and depends on electron flow to PSI.

To further explore the role of PSI in generating the LDT oxidative burst, repetitive light pulses (rSP) were employed to induce PSI photoinhibition (Sejima et al., 2014; Zivcak et al., 2015); rSP of 15,000 µmol m² s⁻¹ was applied every 10 s for 15 min, followed by 9 min of recovery (Fig. 4a). This treatment resulted in a decrease of ΦPSI from 0.96 to 0.55 and of ΦPSII from 0.81 to 0.63, accompanied by increased NPQ and PSI acceptor side limitation (ΦNA) (Supp. Fig. 8). Ultimately, ΦPSI and ΦPSII values recovered to 66% and 91% of their initial quantum potential, respectively (Fig. 4a). These findings are consistent with the suggestion that rSP specifically photoinactivates PSI (Sejima et al., 2014; Zivcak et al., 2015). To further examine the effect of rSP on electron transfer downstream to PSI, MV-induced roGFP oxidation was assessed in specific regions of the leaf that were subjected to rSP. Areas with active PSI showed a sharp roGFP2 oxidation spike, reflecting MV-driven electron diversion and superoxide formation, while the PSI photoinactivated regions showed minimal oxidation (Fig. 4b, Supp. Fig. 9). These results demonstrate that rSP-induced damage to PSI reduced electron flow to MV and thus diminished MV-induced roGFP2 oxidation. Of note, rSP-treated regions exhibited chl-roGFP2 responses to both DTT and H₂O₂, which were identical to untreated regions, indicating that the rSP protocol does not compromise probe sensitivity (Supp. Fig. 10).

Notably, the area with photoinactivated PSI showed a lower oxidation peak during the transition to darkness, as demonstrated by the lower chl-roGFP2 oxidation state, compared to untreated leaves or the untreated region of the same leaf (mock) (Fig. 4d & e). Similarly, dose-dependent inactivation of ΦPSI (0–80%) resulted in a gradual decline in LDT-derived chl-roGP2 oxidation, with 80% inactivation completely abolishing the response (Fig. 4e). These findings underscore the importance of active PSI integrity for generating an E_GSH_ oxidative burst during LDT.

### PSI photodamage inhibits dark-associated inactivation of carbon assimilation

LDT-induced chl-roGFP2 oxidation was further examined in the *pgr5* mutant, which serves as a model for studying PSI photodamage (Lima-Melo et al., 2019; Wada et al., 2021), as well as in *pgrl1ab* plants, which display a HL-hypersensitive phenotype similar to that observed in *pgr5*. These plants are deficient in PGRL1, a protein required for PGR5 stabilization and activity (Hertle et al., 2013; Rühle et al., 2021; Suorsa et al., 2016). chl-roGFP2 OxD slopes during LTD in both plant lines were 12% and 30% lower compared with WT, respectively. (Fig. 5a and b, Supp Fig. 11). A similar trend showing a lower LDT-associated oxidative peak in the mutant lines, along with a difference in NPQ induction in the mutants compared to WT, was observed using wavelength-resolved fluorescence (Hipsch et al., 2025) (Supp. Fig. 12). For comparison, the OxD slope in NON-PHOTOCHEMICAL QUENCHING 1 mutant plants (*npq1)* was higher than WT (7.5%), while in STATE TRANSITION 7 mutant plants (*stn7*), values were unchanged (Fig. 5b; Supp. Fig. 11). Consistent with earlier works, *npq1*, *pgr5*, and *pgrl1ab* all displayed similarly reduced NPQ, whereas *stn7* showed a pattern comparable to WT (Supp. Fig. 13; (Bellafiore et al., 2005; Hertle et al., 2013; Munekage et al., 2002; Niyogi et al., 1998)). Collectively, these data indicate that photodamaged PSI is associated with decreased *E_GSH_* oxidation during LDT.

Assuming the oxidative peak reflects a buildup of electron acceptors required for CBC enzyme inactivation, it was hypothesized that the weakened LDT-derived oxidation in *pgr5* and *pgrl1ab* would result in a corresponding decrease in carbon assimilation inactivation (Fig. 1a–b). Indeed, measurement of CO_2_ assimilation induction slopes after 1 or 6 min in dark (Fig. 1) showed a 31.5% decrease in activation in WT (6 min compared to 1), while *pgr5* and *pgrl1ab* showed 24.3% and 17.7% decreases (Fig. 5c-d), respectively (p = 0.0023 and 4.3*10^-5^, for *pgr5* and *pgrl1ab* compared to WT, respectively, Fig. 5d ). No changes in inactivation were observed in *npq1* and *stn7* plants (Supp. Fig. 14).

To further explore carbon assimilation inactivation in the *pgr5* mutant, photosynthesis induction slopes were compared in dark-adapted WT and *pgr5* plants exposed to ambient air or N₂. N₂ exposure was applied either throughout the entire experiment (Supp. Fig. 15) or restricted to the initial light phase and the transition to darkness (N_2_-LDT, Fig. 5e, see Methods). Comparisons of plants exposed to air with those treated with N₂ during the LDT and shifted back to ambient air in darkness, revealed a 28% inactivation in WT, whereas *pgr5* mutants showed only 14% inactivation (p-value = 2.2 * 10⁻⁶; Fig. 5e–g). A similar trend was observed in WT and *pgr5* plants exposed to N₂ throughout the experiment (Supplementary Fig. 15). Collectively, these results demonstrate the role of PSI-dependent O₂ reduction and *E_GSH_* oxidation in the inactivation of carbon assimilation enzymes.

## Discussion

The reductive activation of carbon assimilation proteins during photosynthetic induction requires the buildup of reducing equivalents derived from absorbed solar energy, which are converted via the photosynthetic electron transport chain and transmitted to target proteins through the redox regulatory network. Accordingly, oxidative inactivation during photosynthesis cessation would require the buildup of electron acceptors, in the form of H₂O₂, that serve as a sink for reducing power. The data presented here provide experimental and physiological evidence for a mechanism in which a burst of H₂O₂ is generated through the Mehler reaction at PSI during the transition from light to dark. This burst, reflected in the oxidation of chloroplastic *E_GSH_,* serves as an electron acceptor that drives the oxidative inactivation of photosynthetic proteins. This model is supported by the suppression of carbon assimilation inactivation in the absence of oxygen (Fig. 1), the oxygen-dependent *E_GSH_* oxidation burst observed upon transition to darkness (Fig. 2), and the attenuation of both *E_GSH_* oxidation and enzyme inactivation when PSI activity is inhibited (Figs. 3-5). Collectively, these findings position PSI-driven O₂ reduction via the WWC as the primary process underlying dark-induced inactivation of carbon assimilation.

The oxygen dependence of the chloroplastic *E_GSH_* oxidation burst indicates that it originates from H₂O₂ production through the WWC rather than from a decline in NADPH or reduced glutathione reductase activity. Assuming WWC flux is constant in steady light, it remains unclear what drives the oxidative burst upon an abrupt onset of darkness. The burst may arise from a shift in the delicate balance between WWC-driven H₂O₂ production at PSI and its removal by the antioxidant system during the transition to darkness. Accordingly, under constant illumination, H₂O₂ levels are maintained at a steady state by a balance between continuous production and removal. When darkness is imposed, NADPH synthesis ceases, while metabolic pathways continue to consume NADPH at a high rate, leading to a rapid decline in its levels (Lim et al., 2020). The depletion of NADPH suppresses antioxidant capacity, leading to a transient accumulation of H₂O₂. Such a temporal imbalance may create a dynamic window during which H₂O₂ can be redirected to serve as an electron acceptor, driving the oxidative inactivation of redox-regulated proteins. Thus, transient changes in the balance between H₂O₂ generation and its scavenging by the GSH–ascorbate cycle determine the residual pool of oxidizing equivalents available for the oxidative inactivation pathway.

The kinetics and magnitude of the oxidative burst are governed by the light environment preceding darkness (Fig. 2h-k). The kinetics of the chl-roGFP2 oxidative burst likely reflect the accumulation of H₂O₂ and its reduction by the electron flux derived from redox-regulated proteins during the dark phase. These two opposing processes may account for the bell-shaped pattern of chl-roGFP2 oxidation observed during LDT (Fig. 2i). More specifically, at LL, H₂O₂ production is low, resulting in a minimal effect on chloroplastic *E_GSH_*, while at HL, the combination of high H₂O₂ levels and the strong reducing state of redox-regulated proteins results in a burst that is rapidly quenched to oxidation levels lower than those measured under illumination.

Exploring chloroplastic *E_GSH_* redox dynamics under diverse physiological conditions revealed that LDT elicits one of the most pronounced oxidative responses detected thus far with chl-roGFP2 (Haber et al., 2021; Hipsch et al., 2021, 2023). Quantitatively, the roGFP2 redox potential shifted by 10–15 mV under HL conditions, whereas even severe water stress induced only a modest 6.5 mV increase (Hipsch et al., 2021), while the LDT in non-stressed plants caused a notable shift of 10–20 mV (Fig. 2). The observation that chloroplastic *E_GSH_* undergoes reversible shifts between two oxidative states under normal conditions suggests that these shifts reflect a regulatory reconfiguration of metabolic activity through redox control, rather than uncontrolled ROS production merely being scavenged by the GSH pool. Accordingly, alterations in chloroplastic *E_GSH_* observed under stress conditions may represent metabolic adaptation rather than uncontrolled oxidative stress.

The presented oxygen depletion experiments demonstrated that roGFP2 oxidation during LDT was strongly attenuated under low O₂, leaving only a minor residual signal compared to ambient conditions, linking *E_GSH_* oxidation to WWC activity (Fig. 2). Chloroplastic *E_GSH_* oxidation may not merely mirror ongoing changes in H_2_O_2_ levels but may also directly participate in the oxidative inactivation of photosynthetic proteins. The accumulation of oxidized glutathione (GSSG) may function as an electron acceptor in the oxidation of redox-regulated chloroplast proteins, contributing to their dark-induced oxidative inactivation. In this proposed oxidative route, glutaredoxins (GRXs), which catalyze thiol–disulfide exchange reactions between glutathione (GSH) and target proteins, could mediate oxidative inactivation, providing an alternative to the established 2-Cys peroxiredoxin/atypical thioredoxin pathway. This possibility is further supported by the potential GRX activity of thioredoxin-like proteins (TrxL) (Chibani et al., 2012; Jacquot, 2018), a key component of the chloroplast oxidative pathway (Yoshida et al., 2018). Future investigations, particularly those employing GRX-deficient mutant lines in which the redox state of target proteins is decoupled from the *E_GSH_* (Bohle, Rossi, et al., 2024), are required to elucidate the role of the GSH/GRX pathway in the oxidation of stromal proteins during the transition to darkness.

The *pgr5* knockout plants are defective in the induction of photosynthetic control and NPQ, rendering them vulnerable to high and fluctuating light conditions (Munekage et al., 2002; Suorsa et al., 2012; Tikkanen et al., 2010; Yamamoto & Shikanai, 2019). Specifically, damage to the Fe–S clusters of PSI has been demonstrated in *pgr5* mutants under HL, establishing *pgr5* as a valuable genetic model for investigating PSI photodamage (Tiwari et al., 2016, 2024). In the presented experiments, *pgr5* plants exhibited a lower oxidative burst during LDT (Fig.5, a-b). These results are consistent with the misregulation of chloroplastic *E_GSH_* observed in *pgr5* mutants under fluctuating light conditions, in which the mutant plants failed to exhibit the characteristic oscillation between relatively reduced and oxidized states of chl-roGFP2 (Haber et al., 2021). An even more pronounced inhibition of chl-roGFP2 oxidation during LDT was found in *pgrl1ab*. In both mutants carbon assimilation inactivation was also attenuated compared to the WT plant (Fig. 5d). The disrupted redox control and impaired inactivation of the carbon assimilation process in *pgr5* and *pgrl1ab* during LDT may result from the exacerbated damage to the Fe–S clusters in their PSI, as elevated levels of photoinitiated PSI may reduce the capacity for superoxide production via the Mehler reaction. Indeed, markedly attenuated LDT oxidation was also observed in plants treated with rSP (Fig. 4), which has been shown to inhibit the PSI A-branch under HL irradiance, which is the principal site for O_2_ (Kozuleva et al., 2021; Shimakawa et al., 2024).

However, it should be noted that the observed changes in the chl-roGFP2 oxidative burst and the altered regulation of photosynthetic protein inactivation were detected under non-stress conditions, which are not typically associated with PSI photodamage (Tiwari et al., 2016, 2024). While it remains possible that even minor or undetectable damage to PSI Fe–S clusters could influence the Mehler reaction, these results may also point to a direct role of the PGR5/PGRL complex in modulating electron flux through the WWC, thereby positioning it as a key regulator of the oxidative inactivation process. A role for PGR5 in promoting oxidative inactivation of photosynthetic proteins may explain its involvement in downregulation of ATP synthase activity and lowered proton conductivity (gH^+^) during transitions to HL, a redox-dependent regulatory feature conserved across multiple photosynthetic lineages (Nikkanen et al., 2024). As gH⁺ regulation is critical for proper activation of NPQ and photosynthetic control, key photoprotective mechanisms, it is possible that impaired photoprotection and stunted growth observed in *pgr5* mutants under high or fluctuating light result from impaired redox regulation. Additional experiments will be needed to clarify whether the attenuated oxidative burst and diminished inactivation of carbon assimilation observed in *pgr5* and *pgrl1ab* arise solely as consequences of PSI photoinhibition or whether they reflect a direct involvement of the PGR5/PGRL1 complex in the regulation of WWC activity.

An oxidative burst also accompanied the transition from dark to light (Haber et al., 2021), during which the activity of the WWC has been proposed to dissipate excess excitation energy when the photosynthetic electron transport chain is not yet fully operational, generating pH across thylakoid membranes for NPQ formation (Makino et al., 2002; Schreiber & Klughammer, 2016). Recently, oxidation of many redox-regulatory sites has been demonstrated during photosynthetic induction, suggesting that two opposing signals, reductive and oxidative, are associated with the onset of photosynthesis (Doron et al., 2025). Among the oxidized redox-regulated proteins were enzymes associated with nocturnal metabolism, including those involved in starch degradation and chloroplastic glycolysis. Similar to its role during LDT, dawn-associated oxidative burst may contribute to the inactivation of nocturnal enzymes by generating a temporal window that permits oxidative inactivation of redox-regulated proteins. According to this model, the two oxidative peaks observed during dark-to-light and light-to-dark transitions facilitate the rotational shift between autotrophic and heterotrophic modes of plant metabolism.

To conclude, the presented data provide experimental evidence for the role of WWC in mediating the oxidative inactivation of carbon assimilation during LDT, serving as a regulatory module that fine-tunes the redox state of key metabolic enzymes. Consequently, WWC activity is likely crucial for optimization of metabolic flexibility in the rapidly changing light environments encountered in nature. The regulatory role of the WWC may account for its conservation through evolution, despite its inherent production of potentially harmful ROS.

## Materials and Methods

### Plant material, growth conditions and experimental setup

Potato (*Solanum tuberosum* cv. *Desiree*) plants, either wild-type or expressing chl-roGFP2, were vegetatively propagated from cuttings or tubers and grown in 4-L pots filled with a commercial growth medium (Green 761; Even Ari, Beit Elazari, Israel). During propagation, plants were treated with the insecticide Confidor™ (Bayer, neonicotinoid), the fungicide azoxystrobin (Amistar™, Adama), and a nutrient solution containing 1 g L⁻¹ of 20-20-20 NPK supplemented with micronutrients. Plants were initially maintained in a controlled-environment greenhouse (Faculty of Agriculture, Rehovot, Israel) and transferred several days before each experiment to a controlled-environment growth chamber for acclimation. Growth conditions were set to 60–70% relative humidity, ambient CO₂ and a 16-h light/8-h dark cycle with illumination at 120 μmol m⁻² s⁻¹. Unless otherwise indicated, experiments were performed on 3–4-week-old plants in a FytoScope FS-RI 1600 growth chamber (Photon Systems Instruments, Drásov, Czech Republic).

*Arabidopsis thaliana* WT (ecotype Columbia-0), *npq1* (CS3771, At1g08550, obtained from ABRC), *pgr5* (EMS mutant line, At2g05620, obtained from Prof. T. Shikanai), *stn7* (T-DNA insertion line At1g68830, SALK_073254) and *pgrl1ab* lines (Chaturvedi et al., 2024; Haber et al., 2021; Wolf et al., 2020) were used throughout this research. Arabidopsis *pgr5*, *npq1*, and *stn7* mutant lines expressing chl-roGFP2 were generated as previously reported (Haber et al., 2021). To generate the chl-roGFP2-*pgrl1ab* lines, the chl-roGFP2 coding sequence, fused to the first 74 amino acids of PRXa as a chloroplast-targeting signal, was cloned under the CaMV 35S promoter in pART7 and transferred into the binary vector pART27. The construct was introduced into Agrobacterium tumefaciens (GV3101) and transformed into Arabidopsis thaliana by floral dip (Clough & Bent, 1998). Six independent transgenic lines were selected based on chl-roGFP2 fluorescence relative to the WT autofluorescence. Plants were sown on soil (Green 761; Even Ari, Beit Elazari, Israel), placed in 4°C for 2 days to ensure uniform germination, and then grown under 16-h/8-h light/dark cycles with a photosynthetic photon flux density of 120 µmol m^−2^ s^−1^ (21°C, 60%–70% RH, ambient CO_2_) for 2–3 weeks.

### Whole-plant redox imaging

Whole-plant fluorescence was imaged with an Advanced Molecular Imager H.T. (Spectral Ami-HT, Spectral Instruments Imaging, LLC, USA) using Aura software. Imaging setup, calculations and data analysis were performed using established protocols for detached leaves, whole Arabidopsis and potato plants, and leaf discs (Hipsch et al., 2025; Hipsch et al., 2023). Ratiometric images were generated by dividing the corrected 405 nm image by the corrected 465 nm image pixel-by-pixel and displaying the ratios in false color. Pre-processing was performed with a custom MATLAB script.

### Confocal microscopy

Images were acquired using a Leica TCS SP8 Lightning confocal microscope (Leica Microsystems, Wetzlar, Germany) operated with LAS X software and equipped with an HC PL APO CS2 40×/1.10 water-immersion objective. Images were recorded at 2048 × 2048-pixel resolution with a pixel size of 0.142 µm. Chl-roGFP2 fluorescence was excited at 488 nm (15–18% laser power), and emission was collected using a 507–534 nm spectral detection window. Chlorophyll autofluorescence was detected in the 653–732 nm range using the same 488 nm excitation. Detector sensitivity for each spectral band was adjusted using PMT or HyD gain settings optimized to avoid saturation. Merged images were generated in LAS X.

### Chemicals and inhibitor stocks

The following inhibitors were purchased from Sigma-Aldrich Israel Ltd. (Rehovot, Israel) and used in this study: 3-(3,4-dichlorophenyl)-1,1-dimethylurea (DCMU; D2425), prepared in 100% DMSO, 2,5-dibromo-6-isopropyl-3-methyl-1,4-benzoquinone (DBMIB; 271993), prepared in absolute ethanol, 2,6-dichloro-1,4-benzoquinone (DCBQ; 431982), prepared in distilled water, 1,1′-methyl viologen (MV), prepared in DW, N, N′-dicyclohexylcarbodiimide (DCCD; 8.02954), prepared in 100% DMSO, antimycin A (AA; A8674), prepared in 100% DMSO, nigericin (NG; N7143), prepared in 100% DMSO, and N-ethylmaleimide (NEM; 04260), prepared in 100% DMSO. For all experiments, stock solutions were freshly prepared, and inhibitor concentrations were selected based on previously reported doses used to achieve complete inhibition of the corresponding electron-transfer processes.

### Chlorophyll fluorescence measurements

Chlorophyll fluorescence was measured using a PAM Imaging System (IMAG-K7 M-Series MAXI, Walz, Germany). Prior to measurements, intact leaves or leaf discs were dark-adapted for 20 min to ensure full relaxation of non-photochemical quenching. The maximum quantum yield of PSII (Fv/Fm) was determined using a weak measuring beam followed by a saturating pulse, in accordance with standard protocols. During induction measurements, actinic illumination was set to 111 μmol photons m⁻² s⁻¹, and saturating pulses were applied every 20 s for a total duration of 5 min. Images were captured and analyzed using ImagingWinGigE software (version 2.56p, Walz).

### P700 redox kinetics measurements

Whole plants or detached leaves were used to assess P700 redox dynamics. Samples were dark-adapted for 20 min before measurement with a Dual-PAM-100 Fluorescence Measuring System (Walz, Germany). P700 was pre-oxidized under continuous far-red (FR) light. To probe electron flow from PSII, short saturating single-turnover (ST, 50 μs) and multiple-turnover (MT, 50 ms) actinic light (AL) pulses were superimposed on the FR illumination. These induced partial re-reduction of P700, following protocols adapted from Fitzpatrick et al. (2022) and Tiwari et al. (2016). For inhibitor and absorption experiments, entire leaflets were incubated for 1 h in either distilled water or the relevant inhibitor solution, in the dark with gentle rotation. Leaves were then gently blotted dry with Kimwipes before being placed in the measuring chamber. For mutant analysis, P700⁺ kinetics were measured in *pgr5*, *pgrl1ab* and WT, indicating impaired p700 oxidation (Supp. Fig. 17).

### PSI inactivation - repetitive saturated pulse (rSP)

PSI inactivation was performed on detached leaves or intact plants using the method developed by the C. Miyake group (Sejima et al., 2014). Saturating pulses (SP) of red light (300 ms, 15,000 μmol photons m⁻² s⁻¹) were applied every 10 s in the absence of AL, Under ambient air conditions, for 15 min. Following this treatment, the tissue was allowed to recover at the instrument for an additional 9 min, during which an SP was applied every 3 min under the same conditions. SP responses were recorded every 3 min by simultaneous measurement of chlorophyll fluorescence and P700 parameters using either a Dual-PAM-100 or Dual-KLAS NIR system (Walz, Germany). For dose-dependent PSI inactivation, samples were maintained under identical conditions for different time intervals until the desired level of inactivation (ΦPSI = 0–80%) was achieved. Typical inactivation times ranged from ∼5 min (low inactivation, 20%) to 40–60 min (≈80% inactivation). After the inactivation treatment, samples were kept in darkness until reillumination (10 min at 120 μmol photons m⁻² s⁻¹) in the growth chamber, and then quickly moved to whole-plant fluorescence measurements.

### Full-spectrum analysis

Time-resolved biosensor and chlorophyll fluorescence signals were measured using a ChloroSpec spectrometer (ChloroSpec B.V., The Netherlands) operated with the ChloroSoft software. The system features three LED sources (405/20 nm, 465/20 nm, 650/20 nm) and two spectrometers that cover a spectral range of 313–885 nm. The red LED was used for actinic light and saturating pulses. To optimize biosensor detection and prevent chlorophyll saturation, the two spectrometers were set at different sensitivity levels. Short-pass filters (460, 475 and 660 nm) were placed in front of the 405, 465 and 650 nm LEDs, respectively, to block excitation bleed-through into the detectors. Before measurements, fresh detached leaves from WT, *pgr5* and *pgrl1ab* plants expressing chl-roGFP2 were cut and dark-adapted for 20 min. The leaves were illuminated at 120 μmol m⁻² s⁻¹ for 10 min to establish a stable light-adapted state and then subjected to a 30-min LDT protocol. For chl-roGFP2 measurements, 405 and 465 nm excitations were delivered as 10 ms pulses in 2000 μmol m⁻² s⁻¹. Emission signals were integrated between 500 and 550 nm using ChloroKin software and custom MATLAB scripts. Reaction-center closure was triggered with a saturating train of short-flash (STF) pulses (300 pulses, 130 μs width, 3 ms spacing) at 60,000 μmol m⁻² s⁻¹ red light.

### Gas exchange measurements

Carbon assimilation rates were measured with a portable LI-6800 gas exchange analyzer (LI-COR, Lincoln, NE, USA). Conditions were set to 21 °C leaf temperature, 420 ppm reference CO₂ and 70% relative humidity. Potato plants were grown in a controlled-environment growth chamber for 4 weeks prior to measurement. Measurements were taken on the fourth or fifth fully expanded leaf from the shoot apex. After clamping the leaf in the LI-6800 chamber, a 2–3-min acclimation period was allowed under 120 μmol photons m⁻² s⁻¹, with CO₂ and humidity levels matching those during measurement. If needed, light intensity was varied (40, 500, or 1200 μmol photons m⁻² s⁻¹), and the instrument was recalibrated using the automatic matching function at each step.

For *Arabidopsis thaliana*, measurements were performed using the LI-6800 small plant chamber (6800-17). Three plants, 3–4 weeks old and prior to flowering, were grown in 65-mm pots (LI-COR 610-09646) and measured simultaneously. The total leaf area was determined from images captured with an Ami HT imager (excitation power, 5%; emission, 670 nm) and analyzed using MATLAB. Assimilation rates were normalized to the calculated total leaf area and expressed on a per-leaf-area basis (multiplied by a factor of 10 to correct for chamber scaling).

For oxygen-depletion experiments, 99.99% N₂ gas was introduced into the LI-6800 chamber while maintaining illumination at 120 μmol photons m⁻² s⁻¹. The fan speed was set to 5000 rpm to ensure proper mixing. N₂ was manually injected 3 min after the start of the measurement and maintained for 5 min (light-to-dark transition) or throughout the entire experiment, as specified in the experimental design. After depletion, the chamber atmosphere was restored to ambient conditions (21% O₂). For redox imaging of samples exposed to N₂, plants were removed from the LI-6800 chamber at defined time points during the LDT protocol and quickly transferred to the AMI-HT imager for roGFP2 redox state analysis.

For the experiments shown in Figure 2 (d-g), oxygen-depleted conditions were generated using a LI-COR 6800 gas-exchange system combined with whole-plant imaging. Intact potato and Arabidopsis plants or detached leaflets were exposed to 120 μmol photons m⁻² s⁻¹, and baseline measurements were acquired under ambient air. Plants were then briefly exposed to ambient conditions before nitrogen (N₂) was introduced 3 min after the start of the measurement to displace O₂, and the low-oxygen atmosphere was maintained during a subsequent dark period of 0–12 min. Gas-exchange and imaging data were collected at 3-minute intervals throughout the protocol.

Net CO₂ assimilation during the first minutes of reillumination (induction) was normalized for each leaf or plant to its own dynamic range (0 = pre-illumination dark baseline; 1 = maximal induction). For each normalized trace, an individual endpoint for slope estimation was identified as the time point at which the rate of change in carbon assimilation stabilized. Activation slopes were then calculated by fitting a first-order linear regression from the onset of reillumination to this endpoint. Data analysis was performed using a custom MATLAB script that applied predefined fractional-change thresholds to identify the stabilization point and thereby determine the fitting window. Gas-exchange variables were recorded every 5 s throughout the measurements.

The relative activation degree of the Calvin–Benson cycle (CBC) was quantified in two complementary ways:

(1) To assess CBC time-dependent induction under light-dark-light conditions, the ratio of the induction slope measured after 6 min of darkness to that measured after 1 min of darkness under ambient or N₂ was calculated and expressed as a percentage

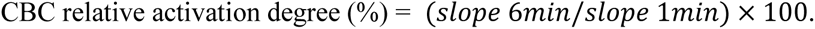

(2) To assess CBC oxygen-dependence induction under light-dark-light conditions, the ratio of the induction slope measured after 6 min of darkness under ambient air to that measured after 6 min of darkness under N₂ LDT was calculated and expressed as a percentage:

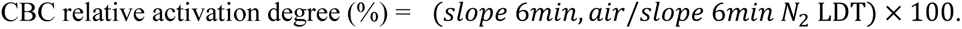

### Statistical and data analysis

Statistical analyses and data visualization were performed using Microsoft Excel 2025, Origin 2025 and custom MATLAB scripts. Differences among treatments were evaluated using one-way ANOVA followed by Tukey–Kramer post hoc tests. When only two groups were compared, statistical significance was assessed using a two-sample, two-tailed Student’s t-test assuming equal variance. Significance was defined as p < 0.05 (*), p < 0.01 (**) and p < 0.001 (***). Box plots were generated using Microsoft Excel’s statistical tools (Microsoft, 2025) or the BoxPlotR web application (Spitzer et al., 2014).

For the box plots, the box indicates the interquartile range (25th–75th percentiles), the horizontal line inside the box represents the median, and the "X" marks the mean. Whiskers extend to display the variability outside the upper and lower quartiles, and individual biological replicates are overlaid as open circles. Significant differences between specific groups are indicated by brackets labelled with asterisks or "n.s." (not significant).

## Supporting information

Supplementary figures_Hipsch et al., 2026

## References

Asada, K. (1999). The water-water cycle in chloroplasts: Scavenging of active oxygens and dissipation of excess photons. Annual Review of Plant Physiology and Plant Molecular Biology, 50(1), 601–639. 10.1146/annurev.arplant.50.1.601

Asada, K. (2000). The water–water cycle as alternative photon and electron sinks. Philosophical Transactions of the Royal Society of London. Series B: Biological Sciences, 355(1402), 1419–1431.

Asada, K., Kiso, K., & Yoshikawa, K. (1974). Univalent reduction of molecular oxygen by spinach chloroplasts on illumination. Journal of Biological Chemistry, 249(7), 2175–2181.

Badger, M. R. (1985). Photosynthetic oxygen exchange.

Badger, M. R., von Caemmerer, S., Ruuska, S., & Nakano, H. (2000). Electron flow to oxygen in higher plants and algae: rates and control of direct photoreduction (Mehler reaction) and rubisco oxygenase. Philosophical Transactions of the Royal Society of London. Series B: Biological Sciences, 355(1402), 1433–1446.

Barry, P., Young, A. J., & Britton, G. (1990). Photodestruction of pigments in higher plants by herbicide action: I. The effect of DCMU (diuron) on isolated chloroplasts. Journal of Experimental Botany, 41(2), 123–129.

Bellafiore, S., Barneche, F., Peltier, G., & Rochaix, J.-D. (2005). State transitions and light adaptation require chloroplast thylakoid protein kinase STN7. Nature, 433(7028), 892–895.

Bohle, F., Klaus, A., Ingelfinger, J., Tegethof, H., Safari, N., Schwarzländer, M., Hochholdinger, F., Hahn, M., Meyer, A. J., & Acosta, I. F. (2024). Contrasting cytosolic glutathione redox dynamics under abiotic and biotic stress in barley as revealed by the biosensor Grx1–roGFP2. Journal of Experimental Botany, 75(8), 2299–2312.

Bohle, F., Rossi, J., Tamanna, S. S., Jansohn, H., Schlosser, M., Reinhardt, F., Brox, A., Bethmann, S., Kopriva, S., & Trentmann, O. (2024). Chloroplasts lacking class I glutaredoxins are functional but show a delayed recovery of protein cysteinyl redox state after oxidative challenge. Redox Biology, 69, 103015.

Buchanan, B. B. (1981). Role of light in the regulation of chloroplast enzymes.

Buchanan, B. B. (2014). The birth of redox regulation. Molecular Plant, 7(1), 1–3.

Buchanan, B. B., & Balmer, Y. (2005). Redox regulation: a broadening horizon. Annu. Rev. Plant Biol., 56(1), 187–220.

Buchanan, B. B., Holmgren, A., Jacquot, J.-P., & Scheibe, R. (2012). Fifty years in the thioredoxin field and a bountiful harvest. Biochimica et Biophysica Acta (BBA)-General Subjects, 1820(11), 1822–1829.

Bus, J. S., & Gibson, J. E. (1984). Paraquat: model for oxidant-initiated toxicity. Environmental Health Perspectives, 55, 37.

Chaturvedi, A. K., Dym, O., Levin, Y., & Fluhr, R. (2024). PGR5-like photosynthetic phenotype 1A redox states alleviate photoinhibition during changes in light intensity. Plant Physiology, 194(2), 1059–1074.

Chibani, K., Tarrago, L., Gualberto, J. M., Wingsle, G., Rey, P., Jacquot, J.-P., & Rouhier, N. (2012). Atypical thioredoxins in poplar: the glutathione-dependent thioredoxin-like 2.1 supports the activity of target enzymes possessing a single redox active cysteine. Plant Physiology, 159(2), 592–605. 10.1104/pp.112.197723

Clough, S. J., & Bent, A. F. (1998). Floral dip: a simplified method for Agrobacterium-mediated transformation of Arabidopsis thaliana. The Plant Journal, 16(6), 735–743.

Dangoor, I., Peled-Zehavi, H., Wittenberg, G., & Danon, A. (2012). A chloroplast light-regulated oxidative sensor for moderate light intensity in Arabidopsis. The Plant Cell, 24(5), 1894–1906.

Dietz, K.-J., Jacob, S., Oelze, M.-L., Laxa, M., Tognetti, V., de Miranda, S. M. N., Baier, M., & Finkemeier, I. (2006). The function of peroxiredoxins in plant organelle redox metabolism. Journal of Experimental Botany, 57(8), 1697–1709.

Doron, S., Lampl, N., Savidor, A., Pri-Or, A., Katina, C., Cejudo, F. J., Levin, Y., & Rosenwasser, S. (2025). Two opposing redox signals mediated by 2-cys peroxiredoxin shape the redox proteome during photosynthetic induction. Redox Biology, 86, 103810. 10.1016/j.redox.2025.103810

Driever, S. M., & Baker, N. R. (2011). The water–water cycle in leaves is not a major alternative electron sink for dissipation of excess excitation energy when CO2 assimilation is restricted. Plant, Cell & Environment, 34(5), 837–846.

Eliyahu, E., Rog, I., Inbal, D., & Danon, A. (2015). ACHT4-driven oxidation of APS1 attenuates starch synthesis under low light intensity in Arabidopsis plants. Proceedings of the National Academy of Sciences, 112(41), 12876–12881.

Exposito-Rodriguez, M., Laissue, P. P., Yvon-Durocher, G., Smirnoff, N., & Mullineaux, P. M. (2017). Photosynthesis-dependent H2O2 transfer from chloroplasts to nuclei provides a high-light signalling mechanism. Nature Communications, 8(1), 49. 10.1038/s41467-017-00074-w

Fantuzzi, A., Allgöwer, F., Baker, H., McGuire, G., Teh, W. K., Gamiz-Hernandez, A. P., Kaila, V. R. I., & Rutherford, A. W. (2022). Bicarbonate-controlled reduction of oxygen by the QA semiquinone in Photosystem II in membranes. Proceedings of the National Academy of Sciences, 119(6), e2116063119.

Foyer, C. H., & Hanke, G. (2022). ROS production and signalling in chloroplasts: cornerstones and evolving concepts. The Plant Journal, 111(3), 642–661.

Foyer, C. H., & Kunert, K. (2024). The ascorbate–glutathione cycle coming of age. Journal of Experimental Botany, 75(9), 2682–2699.

Foyer, C. H., & Noctor, G. (2011). Ascorbate and glutathione: the heart of the redox hub. Plant Physiology, 155(1), 2–18.

Graan, T., & Ort, D. R. (1986). Detection of oxygen-evolving photosystem II centers inactive in plastoquinone reduction. Biochimica et Biophysica Acta (BBA)-Bioenergetics, 852(2–3), 320–330.

Haber, Z., Lampl, N., Meyer, A. J., Zelinger, E., Hipsch, M., & Rosenwasser, S. (2021). Resolving diurnal dynamics of the chloroplastic glutathione redox state in Arabidopsis reveals its photosynthetically-derived oxidation. The Plant Cell. 10.1093/plcell/koab068

Hertle, A. P., Blunder, T., Wunder, T., Pesaresi, P., Pribil, M., Armbruster, U., & Leister, D. (2013). PGRL1 is the elusive ferredoxin-plastoquinone reductase in photosynthetic cyclic electron flow. Molecular Cell, 49(3), 511–523.

Hipsch, M., Lampl, N., Lev, R., & Rosenwasser, S. (2025). Simultaneous recording of biosensors and chlorophyll fluorescence reveals a tight correlation between ETR and oxidative signals. Plant Physiology, 198(2), kiaf222.

Hipsch, M., Lampl, N., Zelinger, E., Barda, O., Waiger, D., & Rosenwasser, S. (2021). Sensing stress responses in potato with whole-plant redox imaging. Plant Physiology, 1–14. 10.1093/plphys/kiab159

Hipsch, M., Michael, Y., Lampl, N., Sapir, O., Cohen, Y., Helman, D., & Rosenwasser, S. (2023). Early detection of late blight in potato by whole-plant redox imaging. The Plant Journal, 113(4), 649–664.

Ilík, P., Pavlovič, A., Kouřil, R., Alboresi, A., Morosinotto, T., Allahverdiyeva, Y., Aro, E., Yamamoto, H., & Shikanai, T. (2017). Alternative electron transport mediated by flavodiiron proteins is operational in organisms from cyanobacteria up to gymnosperms. New Phytologist, 214(3), 967–972.

Jacquot, J.-P. (2018). Dark deactivation of chloroplast enzymes finally comes to light. Proceedings of the National Academy of Sciences, 115(38), 9334–9335. 10.1073/pnas.1814182115

Jahns, P., & Junge, W. (1989). The protonic shortcircuit by DCCD in photosystem II A common feature of all redox transitions of water oxidation. FEBS Letters, 253(1–2), 33–37.

Jiménez-López, J., Casatejada, A., Gálvez-Ramírez, A., Pérez-Ruiz, J. M., & Cejudo, F. J. (2025). Functional relationship of atypical thioredoxins with NADPH-thioredoxin reductase C and 2-Cys peroxiredoxins in Arabidopsis chloroplasts. Journal of Experimental Botany, eraf283.

Klughammer, C., & Schreiber, U. (2016). Deconvolution of ferredoxin, plastocyanin, and P700 transmittance changes in intact leaves with a new type of kinetic LED array spectrophotometer. Photosynthesis Research, 128(2), 195–214. 10.1007/s11120-016-0219-0

König, J., Baier, M., Horling, F., Kahmann, U., Harris, G., Schürmann, P., & Dietz, K.-J. (2002). The plant-specific function of 2-Cys peroxiredoxin-mediated detoxification of peroxides in the redox-hierarchy of photosynthetic electron flux. Proceedings of the National Academy of Sciences, 99(8), 5738–5743.

Kozuleva, M., Petrova, A., Milrad, Y., Semenov, A., Ivanov, B., Redding, K. E., & Yacoby, I. (2021). Phylloquinone is the principal Mehler reaction site within photosystem I in high light. Plant Physiology, 186(4), 1848–1858. 10.1093/plphys/kiab221

Lam, L., Patel-Tupper, D., Lam, H. E., Steen, C. J., Ma, A., Ma, S. A., Leipertz, A., Lee, T.-Y., Fleming, G. R., & Niyogi, K. K. (2024). Resolving an unconventional non-photochemical quenching signature at the light-to-dark transition. BioRxiv, 2010–2024.

Lampl, N., Lev, R., Nissan, I., Gilad, G., Hipsch, M., & Rosenwasser, S. (2022). Systematic monitoring of 2-Cys peroxiredoxin-derived redox signals unveiled its role in attenuating carbon assimilation rate. Proceedings of the National Academy of Sciences, 119(23), e2119719119.

Lee, K. P., & Kim, C. (2024). Photosynthetic ROS and retrograde signaling pathways. New Phytologist, 244(4), 1183–1198. 10.1111/nph.20134

Lim, S.-L., Voon, C. P., Guan, X., Yang, Y., Gardeström, P., & Lim, B. L. (2020). In planta study of photosynthesis and photorespiration using NADPH and NADH/NAD+ fluorescent protein sensors. Nature Communications, 11(1), 1–12.

Lima-Melo, Y., Gollan, P. J., Tikkanen, M., Silveira, J. A. G., & Aro, E. M. (2019). Consequences of photosystem-I damage and repair on photosynthesis and carbon use in Arabidopsis thaliana. Plant Journal, 97(6), 1061–1072. 10.1111/tpj.14177

Makino, A., Miyake, C., & Yokota, A. (2002). physiological functions of the water-water cycle (Mehler Reaction) and the cyclic electron flow around PSI in rice leaves. In Plant Cell Physiol (Vol. 43, Issue 9). https://academic.oup.com/pcp/article/43/9/1017/1823220

Marty, L., Siala, W., Schwarzländer, M., Fricker, M. D., Wirtz, M., Sweetlove, L. J., Meyer, Y., Meyer, A. J., Reichheld, J.-P., & Hell, R. (2009). The NADPH-dependent thioredoxin system constitutes a functional backup for cytosolic glutathione reductase in Arabidopsis. Proceedings of the National Academy of Sciences, 106(22), 9109–9114. 10.1073/pnas.0900206106

Mehler, A. H. (1951). Studies on reactions of illuminated chloroplasts: I. Mechanism of the reduction of oxygen and other hill reagents. Archives of Biochemistry and Biophysics, 33(1), 65–77.

Michalska, J., Zauber, H., Buchanan, B. B., Cejudo, F. J., & Geigenberger, P. (2009). NTRC links built-in thioredoxin to light and sucrose in regulating starch synthesis in chloroplasts and amyloplasts. Proceedings of the National Academy of Sciences, 106(24), 9908–9913.

Miyake, C. (2010). Alternative electron flows (water–water cycle and cyclic electron flow around PSI) in photosynthesis: molecular mechanisms and physiological functions. Plant and Cell Physiology, 51(12), 1951–1963.

Miyake, C., Shinzaki, Y., Nishioka, M., Horiguchi, S., & Tomizawa, K.-I. (2006). Photoinactivation of ascorbate peroxidase in isolated tobacco chloroplasts: Galdieria partita APX maintains the electron flux through the water–water cycle in transplastomic tobacco plants. Plant and Cell Physiology, 47(2), 200–210.

Mizrachi, A., Graff van Creveld, S., Shapiro, O. H., Rosenwasser, S., & Vardi, A. (2019). Light-dependent single-cell heterogeneity in the chloroplast redox state regulates cell fate in a marine diatom. Elife, 8, e47732.

Motohashi, K., Kondoh, A., Stumpp, M. T., & Hisabori, T. (2001). Comprehensive survey of proteins targeted by chloroplast thioredoxin. Proceedings of the National Academy of Sciences, 98(20), 11224–11229.

Müller-Schüssele, S. J. (2024). Chloroplast thiol redox dynamics through the lens of genetically encoded biosensors. Journal of Experimental Botany, 75(17), 5312–5324.

Müller-Schüssele, S. J., Schwarzländer, M., & Meyer, A. J. (2021). Live monitoring of plant redox and energy physiology with genetically encoded biosensors. Plant Physiology, 186(1), 93–109.

Munekage, Y., Hojo, M., Meurer, J., Endo, T., Tasaka, M., & Shikanai, T. (2002). PGR5 is involved in cyclic electron flow around photosystem I and is essential for photoprotection in Arabidopsis. Cell, 110(3), 361–371.

Nikkanen, L., Wey, L. T., Woodford, R., Mustila, H., Kosmützky, D., Ermakova, M., Rintamäki, E., & Allahverdiyeva, Y. (2024). PGR5 is needed for redox-dependent regulation of ATP synthase both in chloroplasts and in cyanobacteria. BioRxiv, 2011–2024.

Niyogi, K. K., Grossman, A. R., & Björkman, O. (1998). Arabidopsis mutants define a central role for the xanthophyll cycle in the regulation of photosynthetic energy conversion. The Plant Cell, 10(7), 1121–1134.

Ojeda, V., Pérez-Ruiz, J. M., & Cejudo, F. J. (2018). 2-Cys peroxiredoxins participate in the oxidation of chloroplast enzymes in the dark. Molecular Plant, 11(11), 1377–1388.

Park, Y.-I., Chow, W. S., Osmond, C. B., & Anderson, J. M. (1996). Electron transport to oxygen mitigates against the photoinactivation of Photosystem II in vivo. Photosynthesis Research, 50(1), 23–32.

Rühle, T., Dann, M., Reiter, B., Schünemann, D., Naranjo, B., Penzler, J.-F., Kleine, T., & Leister, D. (2021). PGRL2 triggers degradation of PGR5 in the absence of PGRL1. Nature Communications, 12(1), 3941.

Sassenrath-Cole, G. F., & Pearcy, R. W. (1994). Regulation of photosynthetic induction state by the magnitude and duration of low light exposure. Plant Physiology, 105(4), 1115–1123.

Schreiber, U., & Neubauer, C. (1990). O2-dependent electron flow, membrane energization and the mechanism of non-photochemical quenching of chlorophyll fluorescence. Photosynthesis Research, 25(3), 279–293.

Schürmann, P., & Buchanan, B. B. (2008). The ferredoxin/thioredoxin system of oxygenic photosynthesis. Antioxidants & Redox Signaling, 10(7), 1235–1274.

Sejima, T., Takagi, D., Fukayama, H., Makino, A., & Miyake, C. (2014). Repetitive short-pulse light mainly inactivates photosystem I in sunflower leaves. Plant and Cell Physiology, 55(6), 1184–1193.

Serrato, A. J., Pérez-Ruiz, J. M., Spínola, M. C., & Cejudo, F. J. (2004). A novel NADPH thioredoxin reductase, localized in the chloroplast, which deficiency causes hypersensitivity to abiotic stress in Arabidopsis thaliana. Journal of Biological Chemistry, 279(42), 43821–43827.

Shavit, N., Dilley, R. A., & San Pietro, A. (1968). Ion translocation in isolated chloroplasts. Uncoupling of photophosphorylation and translocation of K and H ions induced by nigericin. Biochemistry, 7(6), 2356–2363.

Shimakawa, G., Müller, P., Miyake, C., Krieger-Liszkay, A., & Sétif, P. (2024). Photo-oxidative damage of photosystem I by repetitive flashes and chilling stress in cucumber leaves. Biochimica et Biophysica Acta (BBA) - Bioenergetics, 1865(4), 149490. 10.1016/j.bbabio.2024.149490

Souza, P. V. L. (2025). The redox power of thioredoxins: evolution, mechanisms, and redox regulation of carbon–nitrogen metabolism in plants. Discover Plants, 2(1), 94.

Spitzer, M., Wildenhain, J., Rappsilber, J., & Tyers, M. (2014). BoxPlotR: a web tool for generation of box plots. Nature Methods, 11(2), 121–122.

Sun, H., Yang, Y.-J., & Huang, W. (2020). The water-water cycle is more effective in regulating redox state of photosystem I under fluctuating light than cyclic electron transport. Biochimica et Biophysica Acta (BBA)-Bioenergetics, 1861(9), 148235.

Suorsa, M., Järvi, S., Grieco, M., Nurmi, M., Pietrzykowska, M., Rantala, M., Kangasjärvi, S., Paakkarinen, V., Tikkanen, M., Jansson, S., & Aro, E.-M. (2012). PROTON GRADIENT REGULATION5 is essential for proper acclimation of arabidopsis photosystem I to naturally and artificially fluctuating light conditions. The Plant Cell, 24(7), 2934–2948. 10.1105/tpc.112.097162

Suorsa, M., Rossi, F., Tadini, L., Colombo, M., Jahns, P., Kater, M. M., Leister, D., Finazzi, G., Aro, E.-M., & Barbato, R. (2016). PGR5-PGRL1-dependent cyclic electron transport modulates linear electron transport rate in Arabidopsis thaliana. Molecular Plant, 9(2), 271–288.

Tagawa, K., Tsujimoto, H. Y., & Arnon, D. I. (1963). Role of chloroplast ferredoxin in the energy conversion process of photosynthesis. Proceedings of the National Academy of Sciences, 49(4), 567–572.

Terai, Y., Ueno, H., Ogawa, T., Sawa, Y., Miyagi, A., Kawai-Yamada, M., Ishikawa, T., & Maruta, T. (2020). Dehydroascorbate reductases and glutathione set a threshold for high-light–induced ascorbate accumulation. Plant Physiology, 183(1), 112–122.

Tikkanen, M., Grieco, M., Kangasjaݶrvi, S., & Aro, E.-M. (2010). Thylakoid protein phosphorylation in higher plant chloroplasts optimizes electron transfer under fluctuating light. Plant Physiology, 152(2), 723–735. 10.1104/pp.109.150250

Tiwari, A., Mamedov, F., Fitzpatrick, D., Gunell, S., Tikkanen, M., & Aro, E.-M. (2024). Differential FeS cluster photodamage plays a critical role in regulating excess electron flow through photosystem I. Nature Plants, 10(10), 1592–1603.

Tiwari, A., Mamedov, F., Grieco, M., Suorsa, M., Jajoo, A., Styring, S., Tikkanen, M., & Aro, E.-M. (2016). Photodamage of iron–sulphur clusters in photosystem I induces non-photochemical energy dissipation. Nature Plants, 2(4), 1–9.

Trebst, A., Harth, E., & Draber, W. (1970). On a new inhibitor of photosynthetic electron-transport in isolated chloroplasts. Zeitschrift Für Naturforschung B, 25(10), 1157–1159.

Vaseghi, M.-J., Chibani, K., Telman, W., Liebthal, M. F., Gerken, M., Schnitzer, H., Mueller, S. M., & Dietz, K.-J. (2018). The chloroplast 2-cysteine peroxiredoxin functions as thioredoxin oxidase in redox regulation of chloroplast metabolism. Elife, 7, e38194.

Vogel, M. O., Moore, M., König, K., Pecher, P., Alsharafa, K., Lee, J., & Dietz, K.-J. (2014). Fast retrograde signaling in response to high light involves metabolite export, Mitogen-activated protein kinase 6 (MAPK6), and AP2/ERF transcription factors in Arabidopsis. The Plant Cell, 26(3), 1151–1165.

Wada, S., Amako, K., & Miyake, C. (2021). Identification of a novel mutation exacerbated the PSI photoinhibition in pgr5/pgrl1 mutants; caution for overestimation of the phenotypes in Arabidopsis pgr5-1 mutant. Cells, 10(11), 2884.

Wolf, B., Isaacson, T., Tiwari, V., Dangoor, I., Mufkadi, S., & Danon, A. (2020). Redox regulation of PGRL1 at the onset of low light intensity. The Plant Journal, 103(2), 715–725.

Yamamoto, H., & Shikanai, T. (2019). PGR5-dependent cyclic electron flow protects photosystem I under fluctuating light at donor and acceptor sides. Plant Physiology, 179(2), 588–600.

Yoshida, K., Hara, A., Sugiura, K., Fukaya, Y., & Hisabori, T. (2018). Thioredoxin-like2/2-Cys peroxiredoxin redox cascade supports oxidative thiol modulation in chloroplasts. Proceedings of the National Academy of Sciences, 115(35), E8296–E8304.

Yoshida, K., Yokochi, Y., Tanaka, K., & Hisabori, T. (2022). The ferredoxin/thioredoxin pathway constitutes an indispensable redox-signaling cascade for light-dependent reduction of chloroplast stromal proteins. Journal of Biological Chemistry, 298(12), 102650.

Zhang, P., Allahverdiyeva, Y., Eisenhut, M., & Aro, E.-M. (2009). Flavodiiron proteins in oxygenic photosynthetic organisms: photoprotection of photosystem II by Flv2 and Flv4 in Synechocystis sp. PCC 6803. PLoS One, 4(4), e5331.

Zivcak, M., Brestic, M., Kunderlikova, K., Sytar, O., & Allakhverdiev, S. I. (2015). Repetitive light pulse-induced photoinhibition of photosystem I severely affects CO 2 assimilation and photoprotection in wheat leaves. Photosynthesis Research, 126, 449–463.

