## Supplementary figures_Hipsch et al., 2026 for "Dark-induced inactivation of the carbon assimilation process requires a water–water cycle–driven oxidative burst"

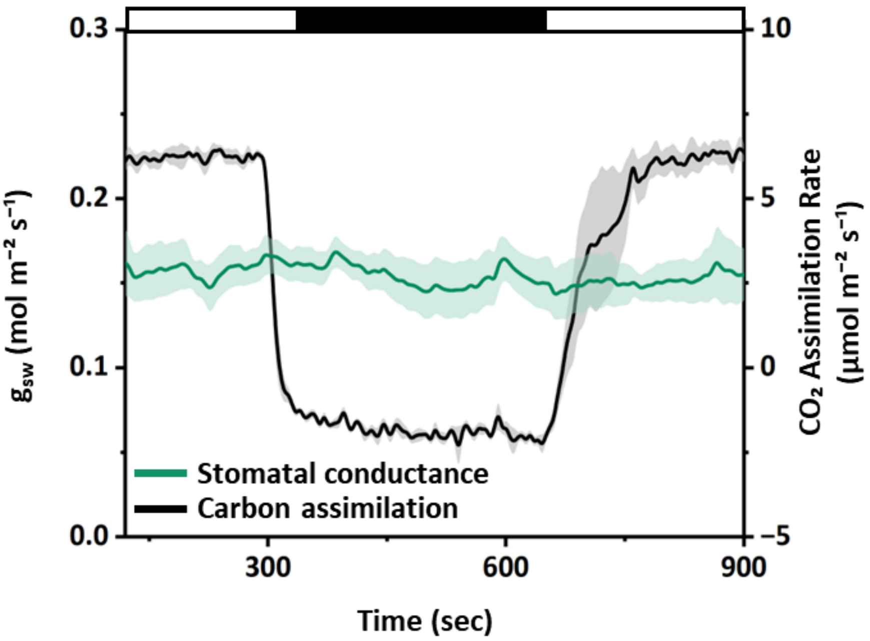


**Supplementary Figure 1: Changes in carbon assimilation and stomatal conductance during light–dark–light transition in potato plants.** The graph shows carbon assimilation and stomatal conductance (gsw) in potato leaves exposed to a light–dark–light schedule of two 5-min illumination periods separated by 6 min of darkness. Data are presented as mean ± SE (n = 4), with shaded areas representing ± SE.

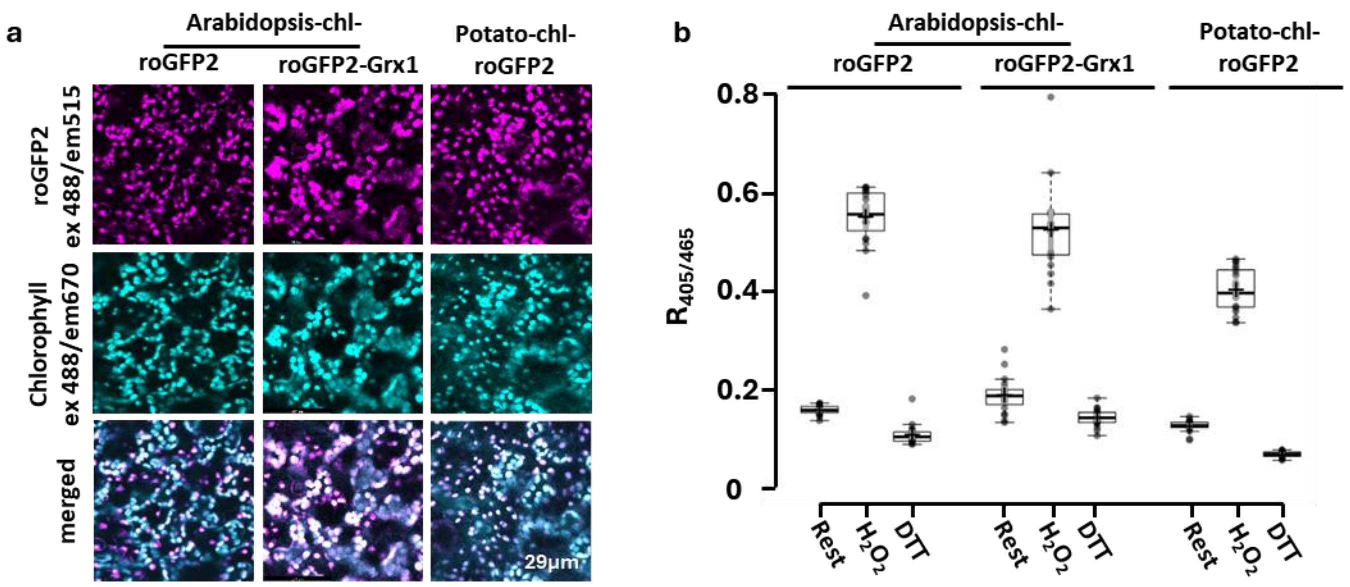


**Supplementary Figure 2: Confocal imaging and calibration of chloroplast-targeted roGFP2 in Arabidopsis and potato.** a) Confocal microscopy images showing chloroplast-targeted roGFP2 (chl-roGFP2) in Arabidopsis and potato, and chloroplast-targeted roGFP2-fused glutaredoxin (Grx1-chl-roGFP2) in Arabidopsis. chl-roGFP2 and chlorophyll autofluorescence were excited at 488 nm. Chl-roGFP2 emission was collected in 507–534 nm, and chlorophyll autofluorescence in 653–732 nm. Merged images combine channels to visualize chloroplast localization. b) Quantification of the biosensors' redox state under resting (DW), oxidized (1 M $H_2O_2$), and reduced (0.1 M DTT) conditions. Arabidopsis plants were infiltrated, and potato leaf discs (0.5 mm²) were submerged in $H_2O_2$ or DTT. Box plots display medians, interquartile ranges (25th–75th percentiles), whiskers (1.5× IQR), outliers (dots), sample means (crosses), and 95% confidence intervals. Data points are presented by open circles (n = 19–20).

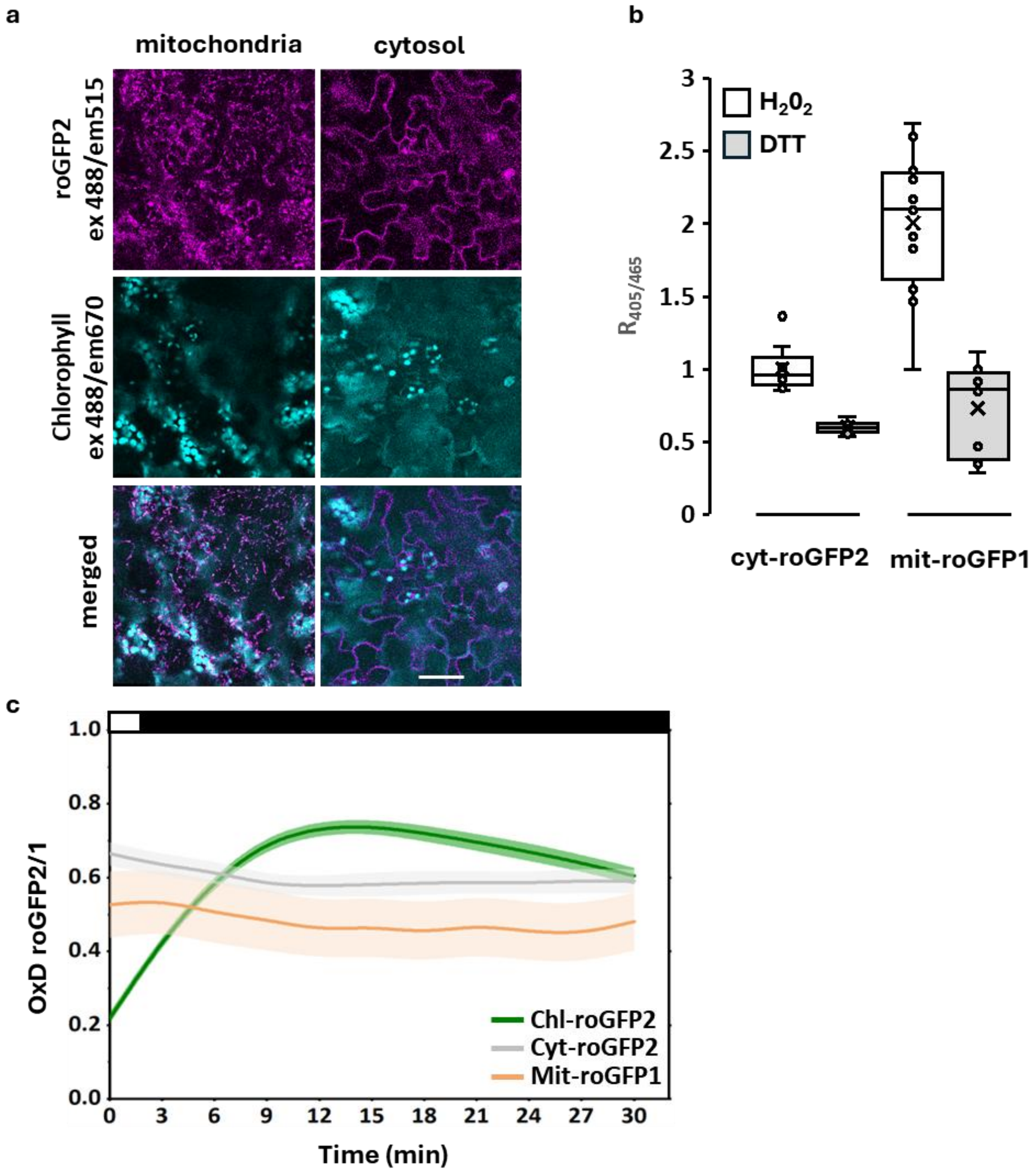


**Supplementary Figure 3**: **chl-roGFP2 oxidation during light-to-dark transitions occurs specifically in chloroplasts and not in the cytoplasm or mitochondria**. a) Confocal microscopy images showing the localization of the roGFP2 biosensors targeted to the chloroplast (chl-roGFP2), mitochondria (mit-roGFP1) and cytosol (cyt-roGFP2). chl-roGFP2 and chlorophyll fluorescence was excited at 488 nm. Chl-roGFP2 emission was collected in 507–534 nm and chlorophyll autofluorescence in 653–732 nm. Merged images combine channels to visualize chloroplast localization. b) Quantification of biosensor redox state under resting (DW), oxidized (1 M $H_2O_2$) and reduced (0.1 M DTT) conditions in cyt-roGFP2 and mit-roGFP1 lines. Data represent mean ± SE (n = 8–11). c) Changes in oxidation degree (OxD) during the light-to-dark transition in Arabidopsis plants expressing chloroplast-, cytosol-, or mitochondria-targeted roGFP sensors. Values represent mean ± SE (n = 12–16), with semi-transparent bands in the same color as the mean line representing ± SE

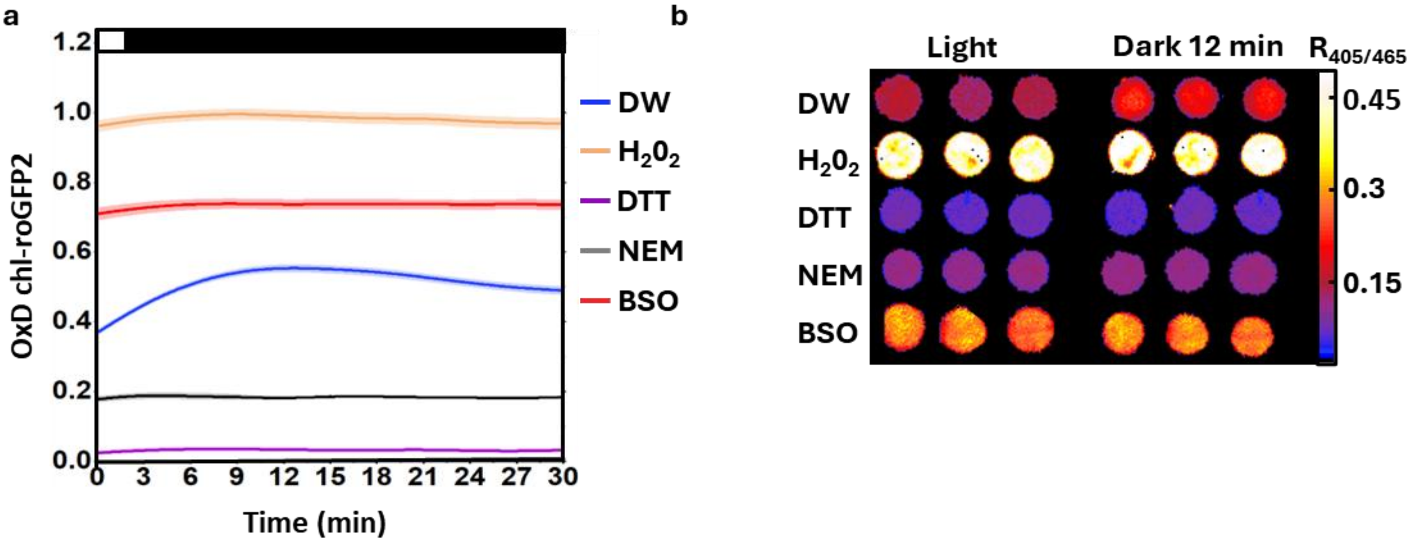


**Supplementary Figure 4**: **Effect of redox-active inhibitors on chl-roGFP2 oxidation during light-to-dark transitions.** a) Potato leaf discs (LD) were submerged in redox-active reagents for 1 h in darkness, followed by 10 min of illumination at 120 µmol $m^{-2}$ $s^{-1}$. Treatments included 1000 mM $H_2O_2$, 100 mM DTT, 20 mM N-ethylmaleimide (NEM) and 2.5 mM L-buthionine- (S, R)-sulfoximine (BSO). Values represent mean ± SE (n = 20–28), with semi-transparent bands in the same color as the mean line representing ± SE. b) Representative ratiometric images of LD submerged in the redox-active reagents listed in (a), for 1 h in darkness.

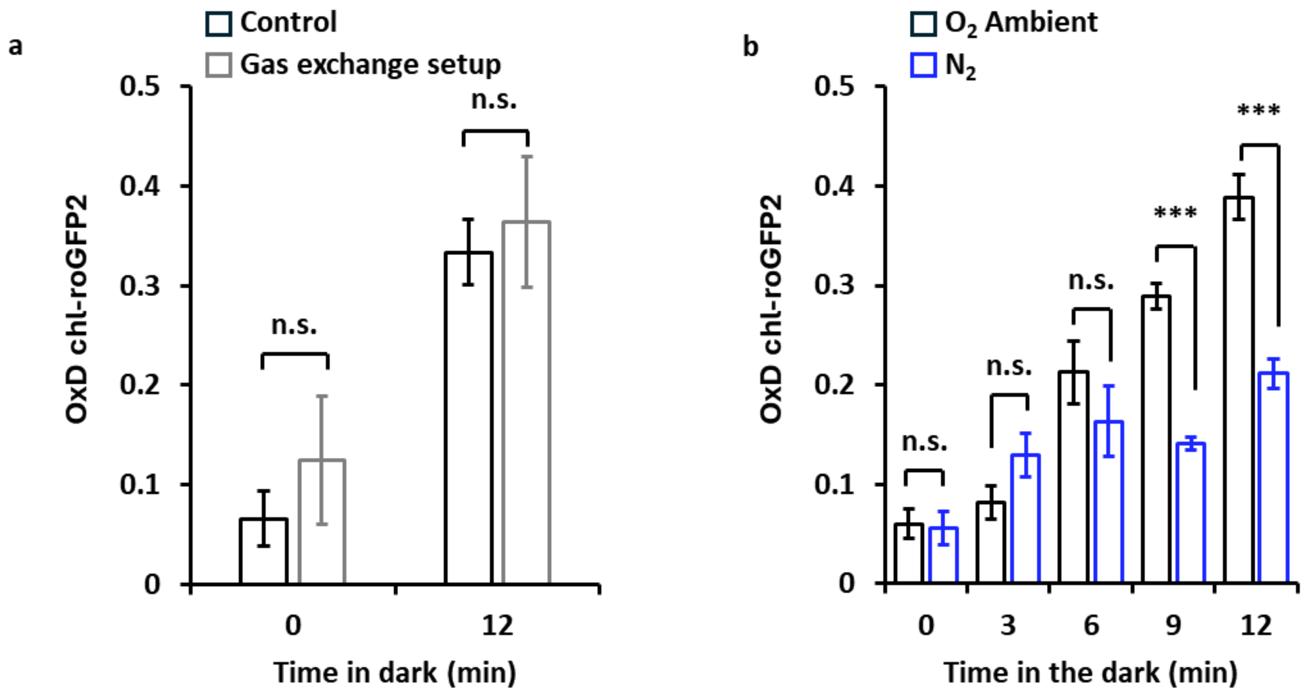


**Supplementary Figure 5: Effect of measurement system and oxygen depletion on chl-roGFP2 oxidation during the light-to-dark transition in potato plants.** a) Comparison of chl-roGFP2 oxidation during LDT in whole potato plants, measured at time 0 (under illumination) and after 12 min of darkness, recorded either in the standard growth chamber or inside the LI-COR 6800 setup. Values represent mean ± SE (n = 3). b) Time-course analysis of chl-roGFP2 oxidation during LDT in whole potato plants exposed to ambient air or to $N_2$ from the onset of darkness (0) until up to 12 min in darkness (0, 3, 6, 9, and 12 minutes). Values represent mean ± SE (n = 5). Statistical significance was assessed for panels (b, c, d, g, and h) using a two-tailed Student's t-test, as is marked as $p < 0.05$ (*), $< 0.01$ (**), $< 0.001$ (***), while "n.s." denotes non-significant differences.

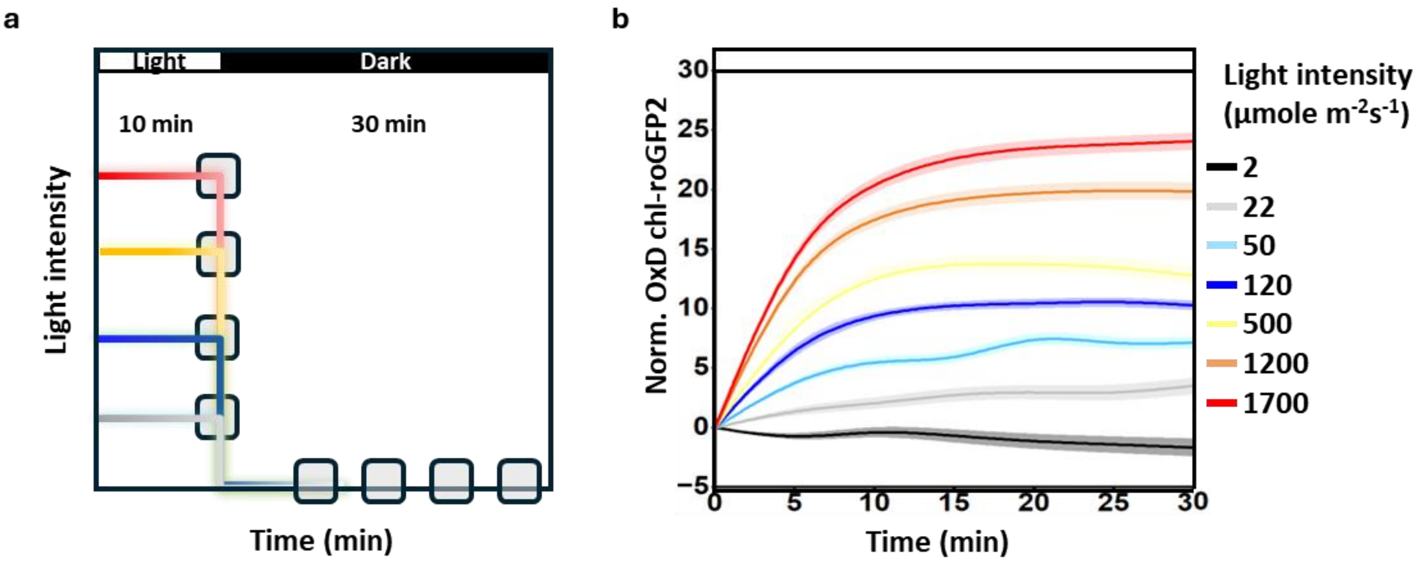


**Supplementary Figure 6: Increasing light intensity enhances oxidation of the chloroplast-targeted roGFP2 probe.** a) Experimental schematic showing the applied dark-to-light conditions. b) Variations in chl-roGFP2 OxD across increasing light intensities (0–1700 µmol $m^{-2}$ $s^{-1}$) during the conditions shown in (a). Values were normalized to chl-roGFP2 OxD in the dark. Detached potato leaf discs (LD, 0.5 $mm^2$) expressing chl-roGFP2 were exposed to the indicated light levels for 30 min, with measurements taken at 5-minute intervals. Values represent mean ± SE (n = 18–21) with semi-transparent bands in the same color as the mean line representing ± SE

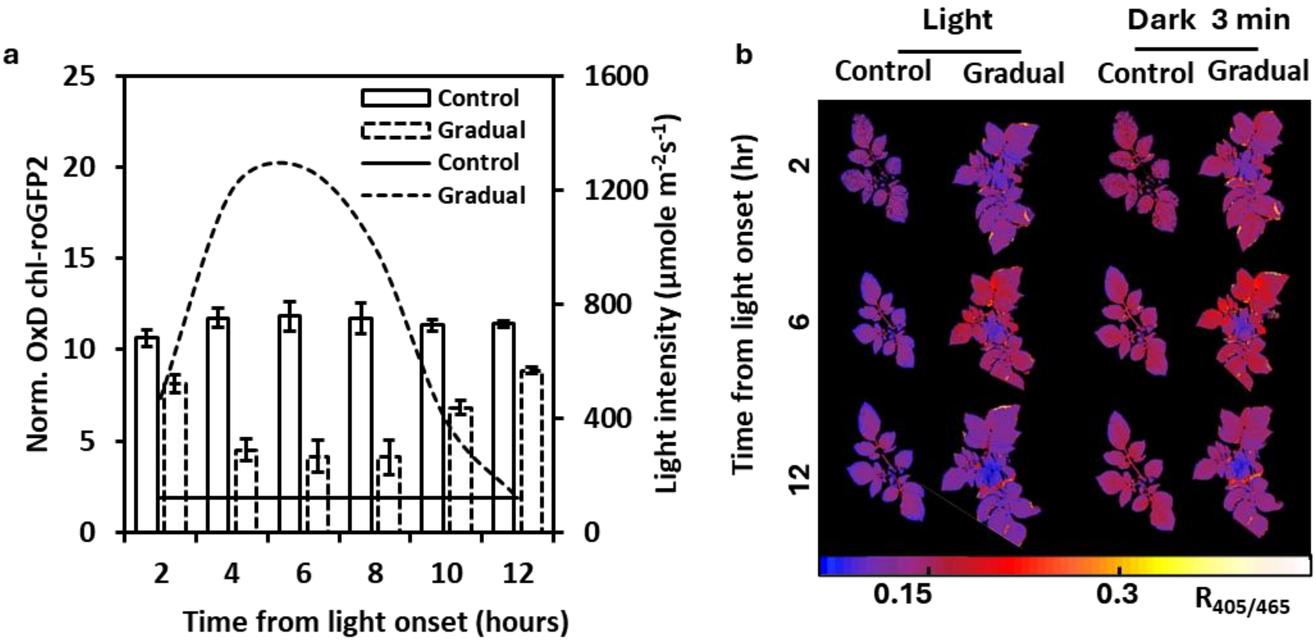


**Supplementary Figure 7: Diurnal changes in chl-roGFP2 oxidation during light-to-dark transitions.** a) Potato plants expressing chl-roGFP2 were exposed to a constant light intensity of 120 µmol $m^{-2}$ $s^{-1}$ throughout the day (control) or to a gradual increase in light intensity reaching up to 1270 µmol $m^{-2}$ $s^{-1}$. chl-roGFP2 oxidation response to LDT was monitored 3 min into the dark phase, and images were collected every two hours. Values represent ± SE (n = 9). b) Representative ratiometric images of potato plants under both control light (120 µmol $m^{-2}$ $s^{-1}$) or a gradual increase in light intensity (up to 1270 µmol $m^{-2}$ $s^{-1}$). Images were captured under light conditions and after 3 min in darkness.

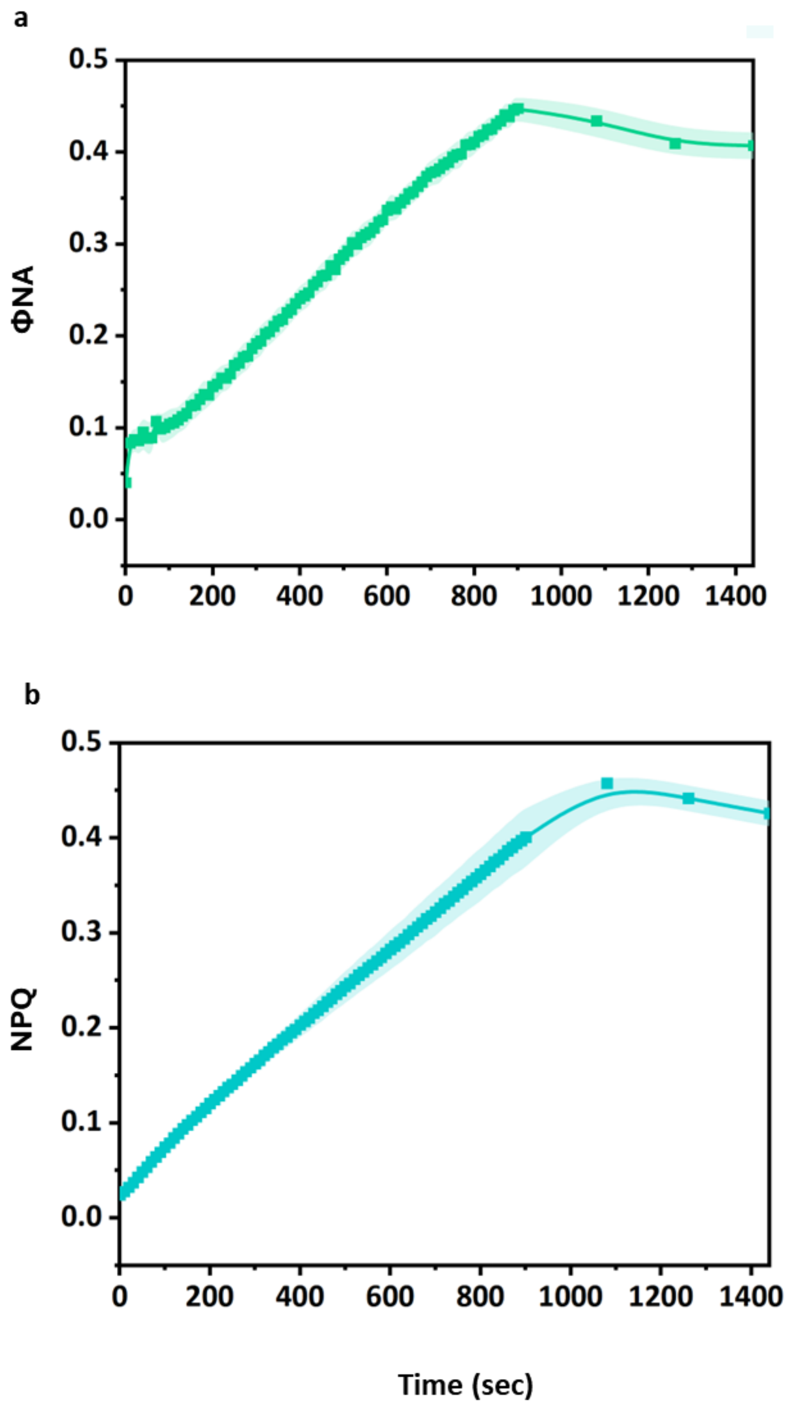


**Supplementary Figure 8: Photosystem I inactivation enhances acceptor-side limitation and non-photochemical quenching.** a) Acceptor-side limitation (ΦNA) and b) non-photochemical quenching (NPQ) in detached leaves exposed to a repetitive saturating pulses (rSP) protocol (See Methods). Measurements were taken during 15 min of rSP, followed by a 9-min recovery phase at 3-min intervals. Data are presented as mean ± SE (n = 4), with semi-transparent bands in the same color as the mean line representing ± SE.

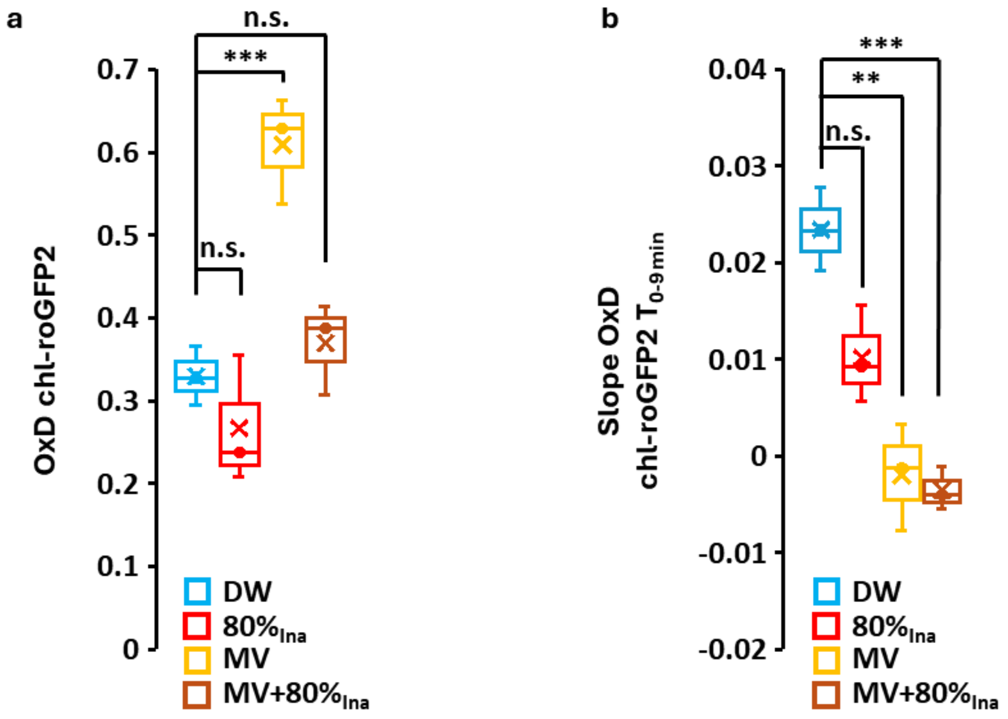


**Supplementary Figure 9: Combined effect of photosystem I inactivation and methyl viologen on chl-roGFP2 oxidation in light and during light-dark transition.** a) Detached leaves were pre-inactivated and subsequently soaked in either 10 µM methyl viologen or distilled water (DW) as a control and then illuminated at 120 µmol $m^{-2}$ $s^{-1}$ for 10 min before imaging. The box plots show changes in chl-roGFP2 OxD in regions of interest (ROIs) exposed to the inactivation process. Values represent mean (n = 4). b) Box plot showing changes in the OxD slope calculated over 0–9 minutes of darkness for the same ROIs. Values represent mean (n = 4). Statistical significance was assessed for panels using a two-tailed Student's t-test and are marked as $p < 0.05$ (*), $< 0.01$ (**), $< 0.001$ (***), while "n.s." denotes non-significant differences.

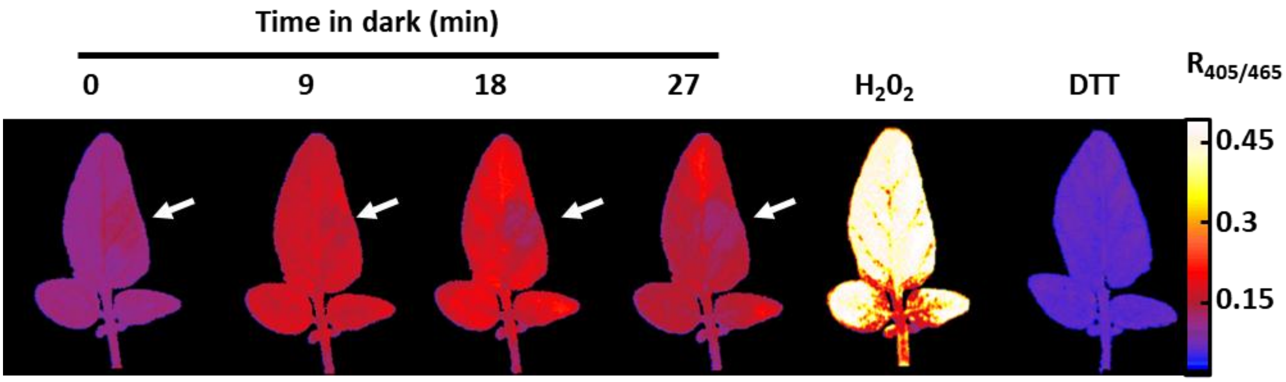


**Supplementary Figure 10: Photosystem I inactivation does not affect chl-roGFP2 redox responsiveness.** Representative ratiometric redox images ($R_{405/465}$) captured during the light-to-dark transition (0–27 min in darkness) in detached leaves subjected to PSI photoinactivation. Following the dark-light transition, leaves were soaked for 10 min in 1000 mM $H_2O_2$, and then for 60 min in 100 mM dithiothreitol (DTT).

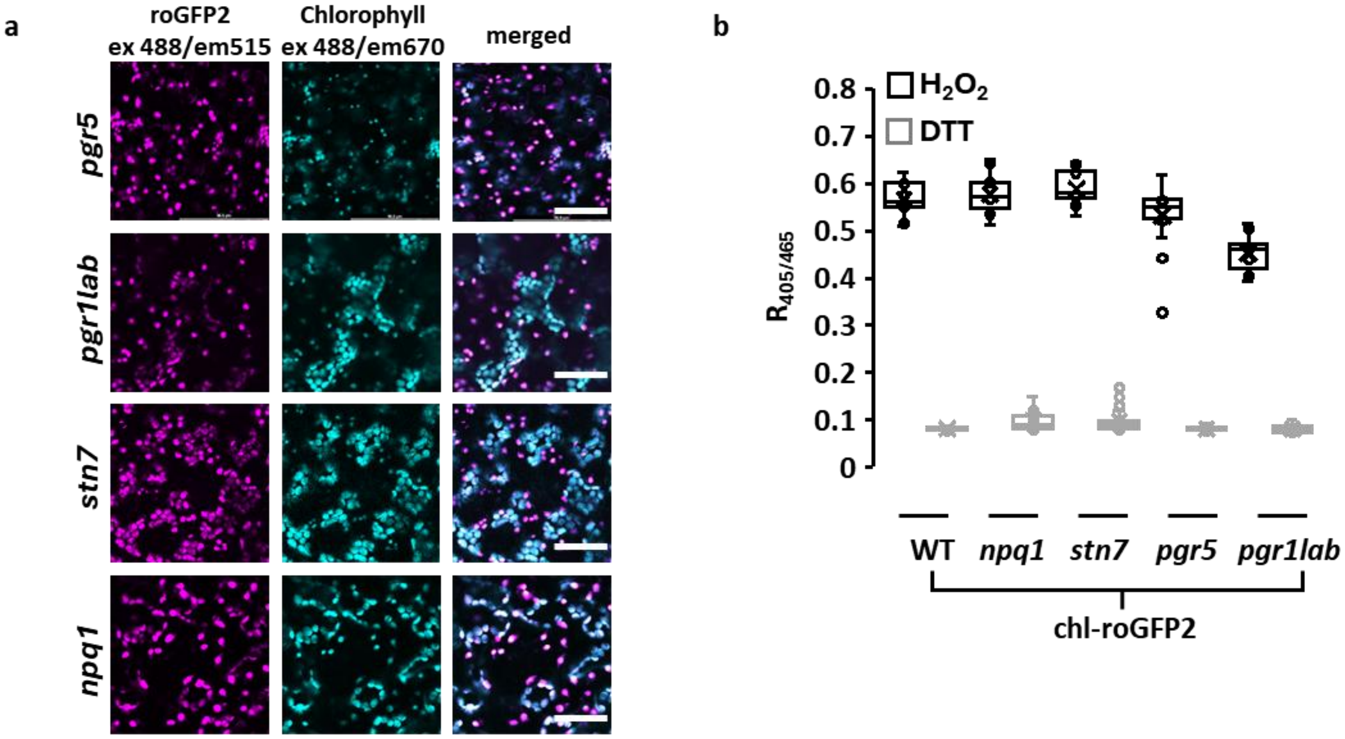


**Supplementary Figure 11: Subcellular localization and redox calibration of chl-roGFP2 in photosynthetic mutants.** a) Confocal microscopy images showing chl-roGFP2 and chlorophyll fluorescence in *pgr5*, *pgrl1ab*, *stn7* and *npq1* Arabidopsis mutants. chl-roGFP2 and chlorophyll autofluorescence were excited at 488 nm. Chl-roGFP2 emission was collected in 507–534 nm and chlorophyll autofluorescence in 653–732 nm. Merged images combine channels to visualize chloroplast localization. b) Box plots showing ratiometric analysis of whole Arabidopsis plants expressing chl-roGFP2 and treated with 1000 mM hydrogen peroxide ($H_2O_2$) or 100 mM dithiothreitol (DTT). Values represent n = 11–20, with means indicated by "X," boxes representing the 25th–75th percentiles and individual samples shown as open circles.

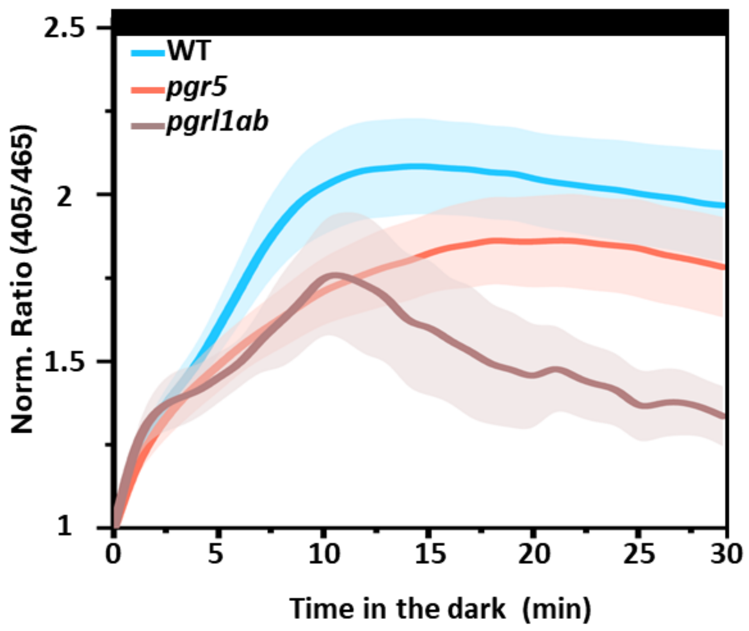


**Supplementary Figure 12: Monitoring chl-roGFP2 oxidation in wild-type, *pgr5* and *pgrl1ab* plants using wavelength-resolved fluorescence data.** Normalized 405/465 ratios recorded during the light-to-dark transition in WT, *pgr5*, and *pgrl1ab* plants expressing chl-roGFP2. Values represent mean ± SE (n = 7–9), with shaded areas indicating ± SE.

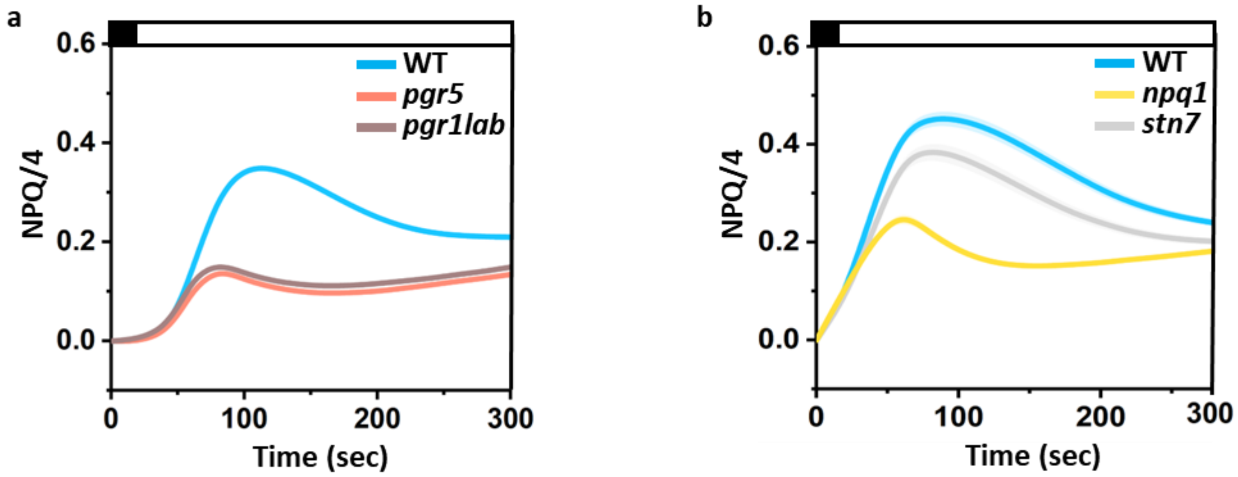


**Supplementary Figure 13: Non-photochemical quenching dynamics in Arabidopsis wild type and photosynthetic mutants.** a) Induction curves of non-photochemical quenching (NPQ/4) in Arabidopsis wild-type (WT) and cyclic electron flow mutants *pgr5* and *pgrl1ab*. Values represent mean ± SE (n = 16), with semi-transparent bands in the same color as the mean line representing ± SE. b) Induction curves of non-photochemical quenching (NPQ) in WT, *npq1* and *stn7* mutants. Values represent mean ± SE (n = 20), with shaded areas indicating ± SE.

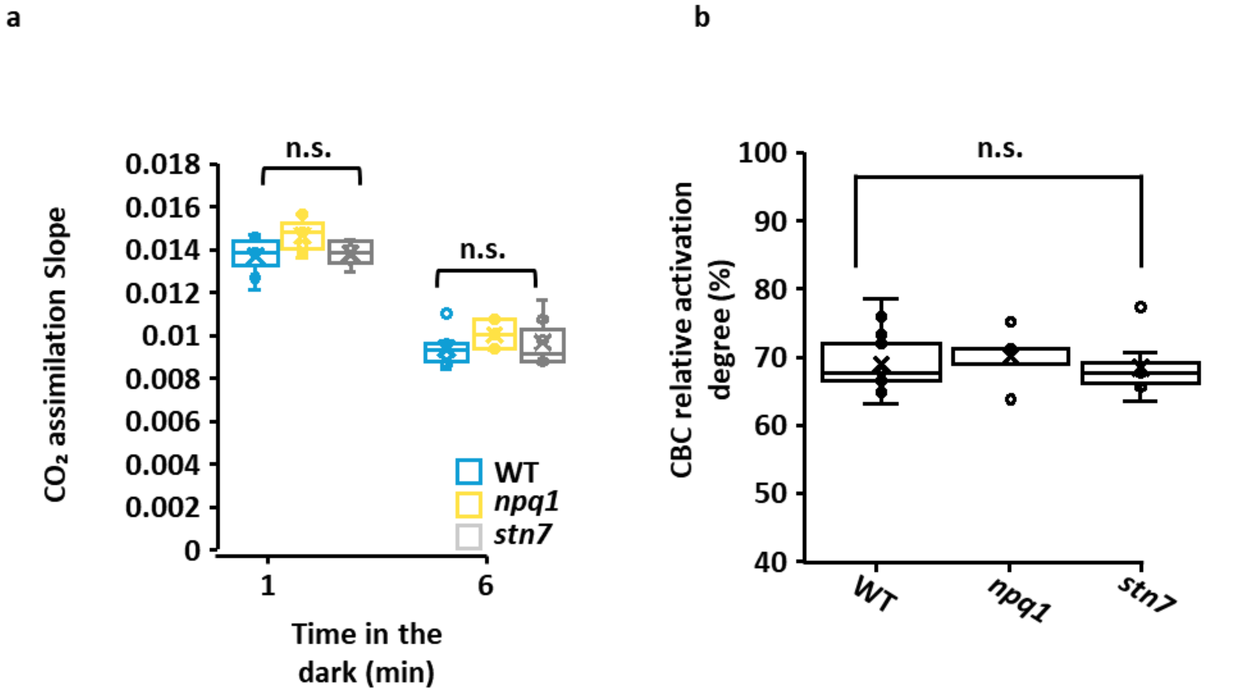


**Supplementary Figure 14: Carbon assimilation induction slopes in photosystem II-related mutants.** a) Box plots showing CO assimilation slopes in Arabidopsis WT, *npq1* and *stn7*, following dark inactivation periods of 1 or 6 minutes. Values are presented as mean ± SE (n = 5-6), with means indicated by "X," boxes representing the 25th–75th percentiles and individual samples shown as open circles. b) Box plots showing the carbon assimilation activation degree of WT, *npq1* and *stn7*, calculated as $slope_{6min}$ / $slope_{1min}$ × 100 (%). Values are presented as mean (n = 5-6). Statistical significance was assessed for both panels using a two-tailed Student's t-test, with asterisks marking significant differences: $p < 0.05$ (*), < 0.01 (**), < 0.001 (***).

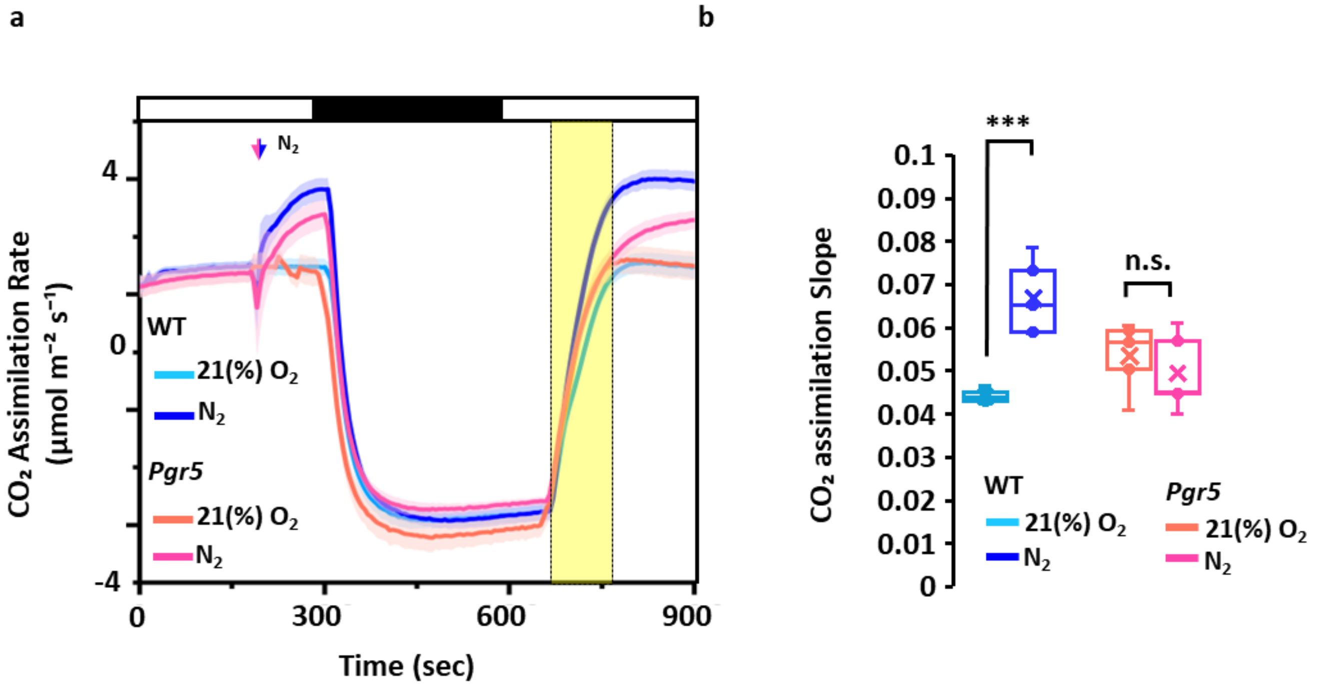


**Supplementary Figure 15: Effect of oxygen availability on carbon assimilation induction slopes during light–dark–light transitions in WT and *pgr5*.** a) Carbon assimilation rates were measured during light-dark-light periods in WT and *pgr5* exposed to 21% $O_2$ or continuous $N_2$ exposure (initiated after 3 min of the first light period). Data are expressed as mean ± SE (n = 4), with shaded regions representing ± SE. b) Box plots showing carbon assimilation induction slopes calculated from the area in (a) marked with yellow. Values are presented as mean ± SE (n = 6–9). Statistical significance was assessed for both panels using a two-tailed Student's t-test and is marked as $p < 0.05$ (*), $< 0.01$ (**), $< 0.001$ (***), while "n.s." denotes non-significant differences.

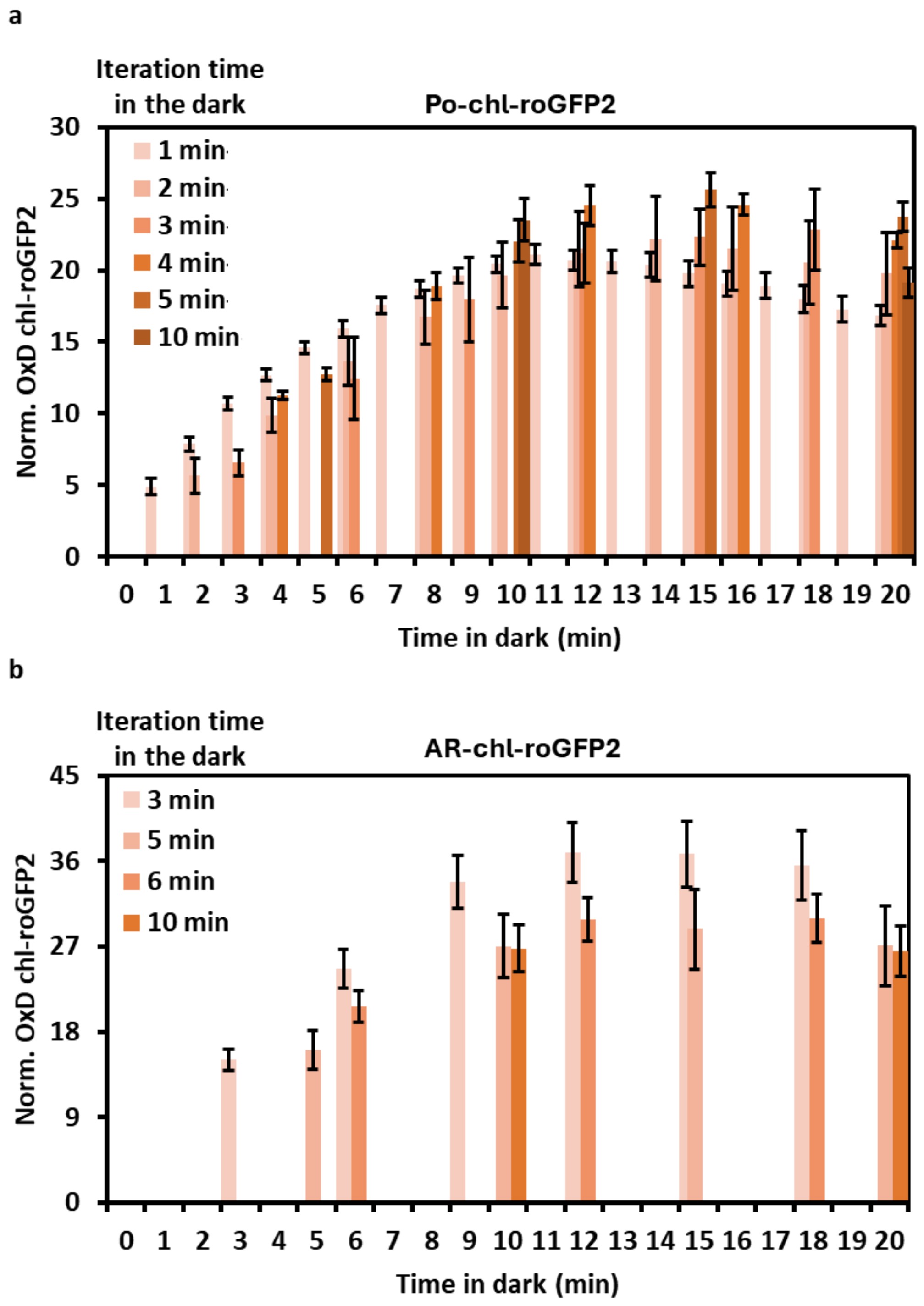


**Supplementary Figure 16: Time-series of chl-roGFP2 oxidation during light-dark transition.** (a) Changes in the chl-roGFP2 OxD in potato (a) and Arabidopsis (b) plants were recorded using a fluorescent camera, with image acquisition intervals between 1 and 10 min during the dark period following illumination (n = 4). Data are presented as mean ± SE. (n=4 and 12 for (a) and (b)).

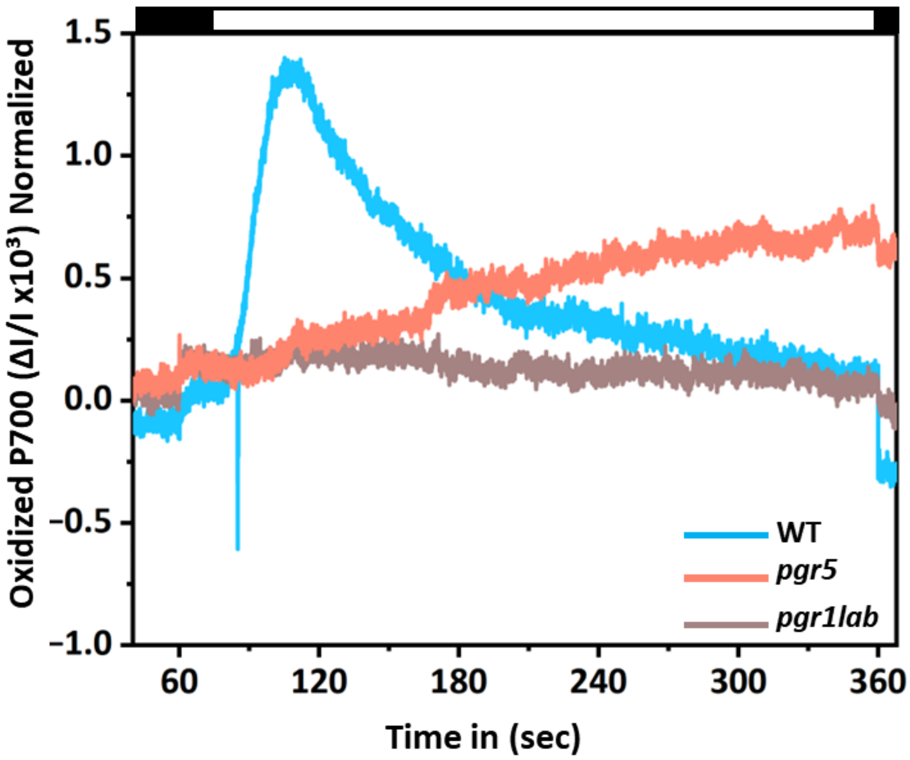


**Supplementary Figure 17: Photosystem I absorbance kinetics in wild type and cyclic electron flow mutants.** Representative P700 redox kinetics in Arabidopsis wild type (WT), *pgr5* and *pgrl1ab* plants. Plants were dark-adapted for 20 min before measurement. Following 30 s in darkness, a saturating pulse (SP) was applied, followed by actinic light (119 µmol $m^{-2}$ $s^{-1}$) for 5 min to reach a steady state. The bars on the top of the graph show a dark-light-dark transition at 0-60 and 300 seconds from the beginning of the experiment. Data represent a single replicate per line; the experiment was repeated three times with similar results.
